# Comparative Chromatin Dynamics of Stem Cell Differentiation in Human and Rat

**DOI:** 10.1101/2021.02.11.430819

**Authors:** Christina Wilcox Thai, Shan Jiang, Yuka Roxas, Cassandra McGill, Savanna Ma, Ali Mortazavi

## Abstract

Differentiation of cell types homologous between species are controlled by conserved networks of regulatory elements driving gene expression. In order to identify conservation of gene expression and chromatin accessibility during cell differentiation in two different species. We collected a daily time-course of gene expression and chromatin accessibility in rat and human to quantify conserved and species-specific chromatin dynamics during embryonic stem cell differentiation to definitive endoderm (DE) as well as to neuronal progenitor cells (NPC). We identify shared and cell-type specific transient differentiation markers in each species, including key transcription factors that may regulate differentiation into each cell-type and their candidate cis-regulatory elements (cCREs). Our analysis shows that DE differentiation has higher conservation of gene expression and chromatin accessibility than NPC differentiation. We provide the first global comparison of transcriptional complexity and chromatin dynamics between human and rat for DE and NPC differentiation.

## INTRODUCTION

A fundamental question in biology is how lineage specification is regulated by a large cohort of transcription factors and signaling molecules. Lineage specification and differentiation are driven, at least in part, by chromatin remodeling causing changes in the accessibility of transcription factor binding sites near promoter and enhancer cis-regulatory modules (CRMs) of other transcription factors (TFs), which collectively form Gene Regulatory Networks (GRNs) (Davidson et al. 2002; Levine and Davidson 2005). Changes in expression of required transcription factors are crucial in the regulation of differentiation. The identification of CRMs that are required for the regulation of gene expression are important for broadening our understanding of development and the conservation or divergence of this development between species and cell lineages (Hardison and Taylor 2012).

Previous functional genomics studies have shown that binding of stage-specific TFs to *cis*-regulatory elements (cREs) controls precise spatiotemporal gene expression. Previous research on *cis-*regulatory element gain or loss (i.e. turnover) has historically been on specific TFs or histone modifications, which limits the observation of species specific or conserved TF-DNA interactions and further prevents the construction of comprehensive GRNs (Berthelot et al. 2018). Furthermore, TF-DNA interactions can be investigated using cRE assays such as Assay for Transposase-Accessible Chromatin using sequencing (ATAC-seq) for TF footprints. TF occupancy near a gene combined with the gene’s expression reveals a regulatory connection which forms the basis of GRNs (F. G. Jørgensen et al. 2005). While individual TF to gene interactions may be conserved weakly, GRNs are highly conserved across species, which makes the comparison of GRNs important for studying evolutionary differences between species (Stergachis et al. 2014).

Embryonic stem cells (ESCs) are defined by their unique ability to self-renew and to generate all lineages of the organism. ESCs divide asymmetrically, producing a daughter cell with more limited differentiation properties (Benitah and Frye 2012). The formation of the three germ layers (ectoderm, mesoderm, and endoderm) in gastrulation is one of the most important developmental processes in the life of an organism as cells lose their indefinite self-renewal capacity. This differentiation process can be mimicked *in vitro* with ESCs (Benitah and Frye 2012). The specification of the germ layers is accomplished through the activation of lineage specific GRNs (Benitah and Frye 2012; Houston 2017). In this study we compare the differentiation of rat and human ES cells into ectodermal neuronal progenitor cells (NPCs) and definitive endoderm (DE) using a time-course of RNA-seq and ATAC-seq in order to determine shared and species-specific evolutionary and regulatory mechanisms that drive development of these two lineages between a common model organism and human.

In the past decade, several groups have established protocols to efficiently differentiate human embryonic stem cells into pancreatic and hepatic cells *in vitro* and to characterize changes in gene expression during the differentiation process (D’Amour et al. 2005, 2006; A. Wang et al. 2015). However, these studies primarily focused on later stages of differentiation and the generation of functional cells, thus leaving the initiation of endoderm formation understudied. Other mammalian species such as mouse and rat have proved surprisingly challenging for robust endoderm layer generation *in vitro* (Kim et al. 2010; Yasunaga et al. 2005; Mfopou et al. 2014). There have been several temporal *in vitro* studies of human ESC to NPC differentiation assessing gene regulatory and cRE dynamics. Some of these include differentiation to cerebral organoids (Kanton et al. 2019), postmitotic neurons (Bunina et al. 2020), early neural differentiation (Shang et al. 2018), primary cultures of human NPCs and the developing brain (de la Torre-Ubieta et al. 2018), and stage-specific transcriptional networks in iPSC neurodevelopment (Siwei Zhang et al. 2018; Forrest et al. 2017). Evolutionary studies have also compared gene expression programs in tissues such as chimpanzee and human comparison (Shang et al. 2018), or mouse and chimpanzee comparisons (de la Torre-Ubieta et al. 2018). Additionally, differences between species in DE differentiation have also been studied (de la Torre-Ubieta et al. 2018c). However, the temporal dynamics of NPC and DE differentiation lineages have not been compared in humans, and the dynamics of rat and human have not been compared for either lineage. Methods for *in vitro* monolayer differentiation of human embryonic stem cells (ESCs) into both definitive endoderm (D’Amour et al. 2005; Kim et al. 2010; D’Amour et al. 2006; Yasunaga et al. 2005; Mfopou et al. 2014; Morrison et al. 2016; Carpentier et al. 2016), and neural progenitor cells (NPCs) (Yap et al. 2015; Muratore et al. 2014; Milani et al. 2016; Lamas et al. 2014; Alsanie et al. 2017) are available. While multiple protocols for mouse differentiation of ESCs to DE have been published (D’Amour et al. 2005; Kim et al. 2010; D’Amour et al. 2006; Yasunaga et al. 2005; Mfopou et al. 2014; Morrison et al. 2016; Carpentier et al. 2016) and NPC (Yap et al. 2015; Muratore et al. 2014; Milani et al. 2016; Lamas et al. 2014; Alsanie et al. 2017), the differentiation of rat ESC into either lineage has not been previously studied using functional genomics.

In order to quantify the extent of conservation of GRNs between rats and humans during lineage specification, we study the differentiation of ES cells into NPC and DE. In order to accomplish this, we first adapted mouse differentiation protocols for rat. We then collected daily time-courses of differentiation for each cell type and species using RNA-seq and ATAC-seq to measure gene expression and chromatin accessibility changes, respectively. We then compared the conservation level of both gene expression and chromatin accessibility in the two species. We compared the RNA-seq and ATAC-seq time-courses between the different lineages in rat, then in human to quantify the dynamics of the transient expressed genes, transcription factor motif enrichment, and to determine their shared function using gene ontology. We then compared human and rat gene modules in each cell type to identify conserved as well as species-specific TF-to-gene connections. Finally, we compared these modules to each other to determine the extent of evolutionarily conserved regulation between the species.

## RESULTS

### Generation of Definitive Endoderm and Neural Progenitor Cells in Two Species

We induced the differentiation of rat and human embryonic stem cells (ESCs) into ectodermal neural progenitor cells and definitive endoderm to quantify the transcriptional and candidate *cis*-regulatory element (cCRE) dynamics that drive cellular commitment. Previously developed methods for *in vitro* monolayer differentiation of human embryonic stem cells (ESCs) into both definitive endoderm (DE) and neural progenitor cells (NPCs) were used (Figure 1A, Figure S1A) (Porterfield 2020; Ramme et al. 2019). We adapted previously published ES mouse differentiation protocols for NPC to rat (Figure 1A, Figure S1A), which have not been previously described (Alsanie et al. 2017; Kim et al. 2010; Yasunaga et al. 2005; Mfopou et al. 2014). For DE differentiation, the protocols for both rat and human rely on the activation of Nodal signaling to induce the expression of TFs and target genes in DE formation. Protocols add exogenous Activin A (Nodal analog) to mimic high concentration of Nodal signaling in order to successfully produce definitive endodermal cells with expression of key marker genes like *MIXL1, CER1, SOX17, FOXA2*, and *CXCR4 (Woodland and Zorn 2008; Teo et al. 2011; Ryan et al. 1996; Conlon et al. 2001; Morrison et al. 2016)*. For NPC differentiation, the protocols for rat and human both rely on Smad inhibition (Porterfield 2020; Alsanie et al. 2017). However, in human NPC differentiation dual Smad inhibition was used, whereas only the inhibitor LDN193189 was used in rat (see methods). Rat NPC differentiation also relies on Fgf2 to further differentiate the cells into a neuronal cell fate (Alsanie et al. 2017; Dwyer 2007). Terminal NPC differentiation was reached by day 8 in both rat and human (Figure 1A and Figure S1A), and terminal DE differentiation was reached by day 7 in rat and day 5 in human (Figure 1B, Figure S1). During each differentiation time-course, morphology changed from a colony like morphology to the formation of neuro-rosettes for NPC differentiation (Figure S1B and 1C) (Fedorova et al. 2019) and pebble/cobblestone morphology of DE differentiation (Figure S1D and 1E) (Siller et al. 2016). We observed morphology changes as early as day 1 of differentiation for both cell-types (Figure S1B-E). Thus the rat cells changed from their ESC morphology to the expected cell type specific morphology over the course of NPC and DE differentiation.

**Figure 1:**
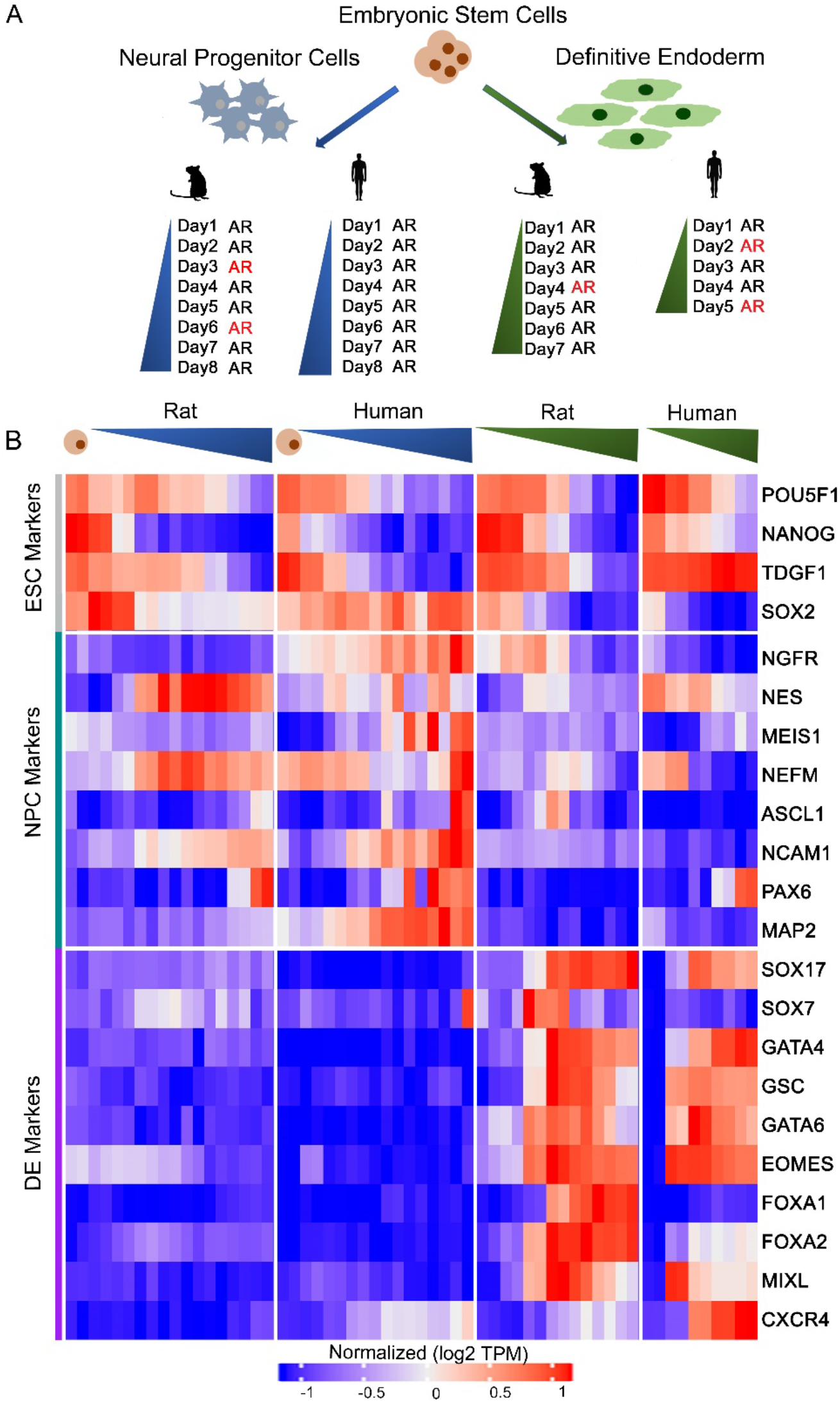
Generation of Definitive Endoderm and Neural Progenitor Cells in Two Species. A) Schematic representation of study design. Color indicates differentiation cell lineage. Human or rat symbols denote species. Letters next to differentiation days (AR) indicate RNA-seq (R) and ATAC-seq (A) timepoints. Red color indicates change in media composition on the specified day. More detailed graphical representation of differentiation outlines including media composition changes in Supplementary Figure 1A. B) Heatmap of marker genes for each differentiation lineage and ESC markers. Color increasing bar on top of the heatmap shows progression of time course. Color indicates differentiation lineage. Cell cartoons indicate ESCs. Colors to the left of the heatmap refer to the lineage groupings the markers belong to. Grey is ESC markers, teal is NPC markers, and purple is DE markers. Transcripts per million (TPM) values were log transformed and row-mean normalized for all genes.

Differentiation of both DE and NPC require the downregulation of key pluripotent genes, *OCT4* and *NANOG* in each species (Figure 1B). However, lingering expression of pluripotency markers have been shown in humans, but does not occur in rodents (Lu et al. 2018; Z. Wang et al. 2012). In order to examine differentiation on a global level for these lineages we checked RNA-seq timepoints daily during differentiation and found that key endodermal markers (*CXCR4, MIXL1, FOXA2, EOMES, GSC, GATA4, SOX17,* and *GATA6*) are activated in both species during definitive endoderm differentiation and key neuronal markers (*NGFR* (Sandberg et al. 2014), *NEFM (Abernathy et al. 2017)*, *NCAM1 (O. S. Jørgensen 1995)*, *NES, PAX6, MAP2, ASCL1 (Tang 2017)*, and *MEIS1* (Owa et al. 2018)) are activated during NPC differentiation (Figure1B). Interestingly, we found that *FOXA1* and *SOX7* were only expressed in rat but not differentially expressed during human DE differentiation. Overall, the increased expression of the lineage specific markers in each species and the decrease in expression of ESC markers suggest successful differentiations into each of the lineages in both species.

### Distinct Transcriptional and Chromatin Accessibility Trajectories for NPC and DE in Rat

We next focused on the comparison of expression and cCRE dynamics between NPC and DE in rat to identify candidate regulators for the differentiation of each cell type. We collected a daily time-course of differentiation in duplicate for RNA-seq and ATAC-seq for a total of 32 RNA-seq and 32 ATAC-seq datasets in rat. We detected 13,112 genes expressed over 1TPM in two or more samples across both NPC and DE differentiation (Table S1). We used maSigPro (Conesa et al. 2006) to identify 10,846 differentially expressed genes that form 18 clusters of distinct expression profiles across time (Figure 2A, Table S2). Five clusters, comprising 2,859 genes (26%), decrease over time in either one or both differentiation time-courses (RR7, RR12, RR14, RR18, RR17). Eight clusters comprising 4,323 genes (40%) increase during NPC differentiation (labeled in blue; RR2, RR4, RR11, RR6, RR15, RR1, RR8, RR16). Five clusters comprising 3,664 genes (34%) increase during DE differentiation (labeled in green; RR3, RR10, RR5, RR9, RR13). We plotted the correlation of the expression profiles over each differentiation time-course (Figure S2A) as well as plotted the samples on UMAP (Figure S2B). We found that UMAP1 separated samples by differentiation time and UMAP2 represents the difference between the two differentiation time-courses. There are larger changes in the differentially expressed genes later in each time-course (Figure S2A and S2B), suggesting a clear switch between early and late differentiation about mid-way through each time-course (Figure S2C). We then focused on the TFs expressed in each of the maSigPro clusters, as they may play a role in the regulation of differentiation into each cell type. As expected, TFs that showed increased expression in NPC clusters include those important in neuronal development and overall expression of early neurons such as *Pax6* (RR11), *Jun* (RR5), *Mef2a* (RR8), *Mef2c* (RR11), *Tead2* (RR1), *Sox9* (RR1), and *bhlhe40* (RR8) (Zhu et al. 2018); (Cho et al. 2011; Dietrich 2013); (“YAP/TAZ Enhance Mammalian Embryonic Neural Stem Cell Characteristics in a Tead-Dependent Manner” 2015)). Similarly, the TFs in clusters that increase during DE differentiation include important markers of DE development and differentiation such as *Foxa2* (RR9), *Eomes* (RR9), *Gata4* (RR9), *Gata6* (RR9), *Foxa1* (RR9), *Lhx1* (RR9), and *Sox17* (RR9). As expected, five clusters with initially high expression that decreased over the time-course (labeled in tan: RR7, RR12, RR14, RR18, RR17) include important ESC TFs such as *Oct4* (RR7), Nrsf/*Rest* (RR7), *Nanog* (RR12), *Sox2* (RR18), and *Tcf7l1* (RR12). The TFs found in cell-type specific clusters in rat suggest that they are expressed in the appropriate pattern to regulate differentiation into NPC and DE.

**Figure 2:**
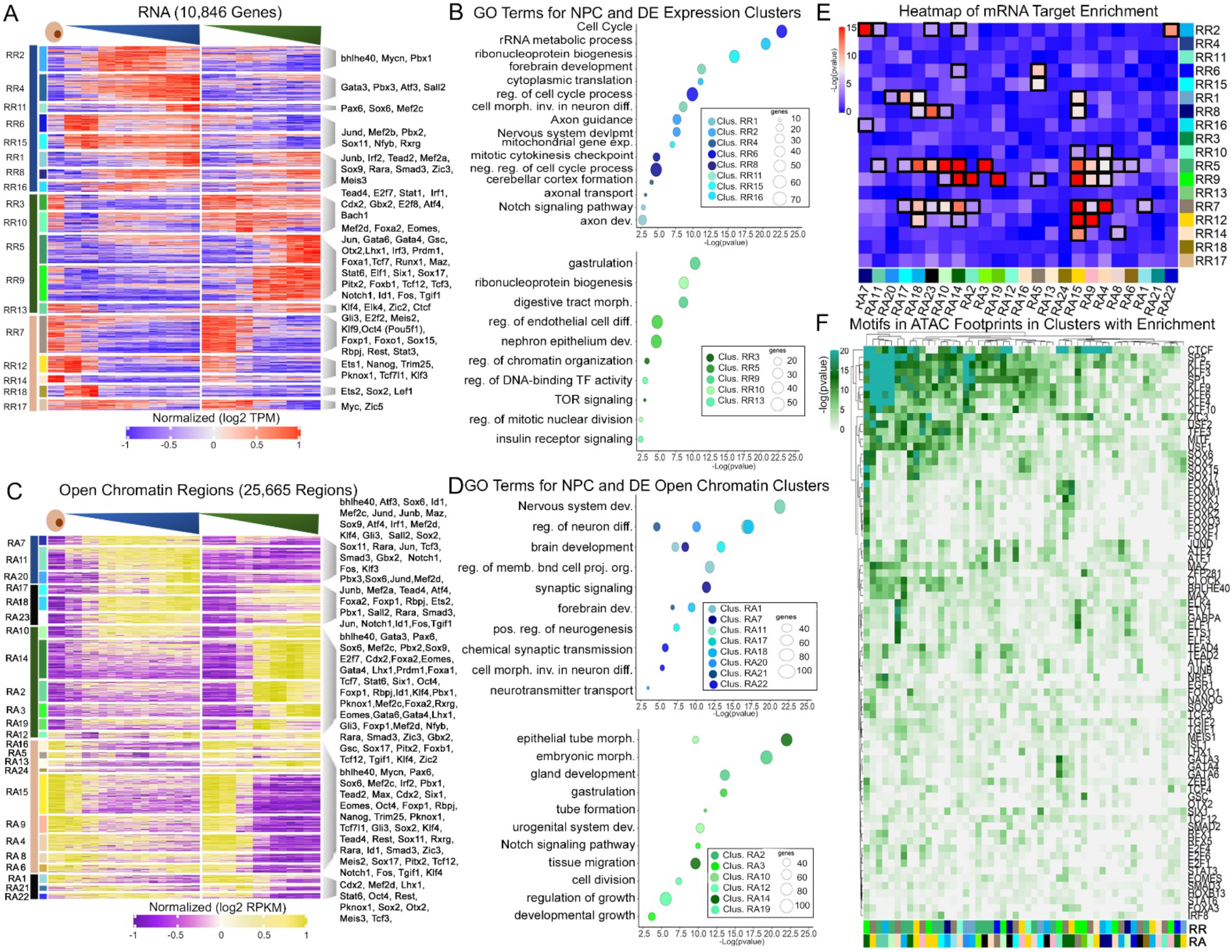
Distinct Transcriptional and Chromatin Accessibility Trajectories for NPC and DE in Rat. A) Heatmap of 10,846 differentially expressed genes (alpha = 0.05, p < 0.05) in NPC and DE differentiation in rat. Hierarchical clustering was used to generate eighteen expression clusters denoted by the adjacent number. Transcripts per million (TPM) values were log transformed and row-mean normalized for all genes. Colors on the far left correspond to the group (NPC, DE, or ESC) the cluster is associated with. The second color bar is for each individual cluster. Select TFs are labeled. B) GO terms for NPC and DE gene expression regions in clusters from Figure 2A. Colors of the clusters correspond to colors next to each cluster in Figure 2A. C) Heatmap of 25,665 differential cCRE regions (alpha = 0.05, FDR = 0.05, p < 0.05) in NPC and DE differentiation in rat. Hierarchical clustering was used to generate twenty four DNA region clusters denoted by the adjacent number. RPKM (Reads Per Kilobase Million) values were log transformed and row-mean normalized for all regions. Regions within 20 kb of the TSS were annotated with the associated gene. Select TFs from Figure 2A were labeled. D) GO terms for NPC and DE cCRE in clusters from Figure 2D. Colors of the clusters correspond to colors next to each cluster in Figure 2C. E) Heatmap of X^2^ p-values on a contingency table of clusters of accessible regions and expression clusters with applied P-value of 10^−4^ (Bonferroni corrected P-value: 0.05/[18 × 24]). The significant cluster overlaps are surrounded by a black box. The colors next to each cluster name match the colors in Figures 2A and 2C. F) Motif enrichments in ATAC footprints for regions with significant overlap between clusters (significant overlaps from Figure 2E). The color bars are the same in A, C, and E for each RNA and ATAC cluster.

We used gene ontology (GO) to analyze clusters higher in NPCs (blue) and DE (green) (Figure 2B) to investigate functional roles of TFs which increased in expression in a cell-type specific manner. Cluster RR11, which sharply increases in expression after day 6 of the NPC time course, is enriched for GO terms such as forebrain development (p < 6×10^−12^) and includes important TFs such as *Pax6* and *Mef2c* (Figure 2B). Cluster RR1, which starts to increase in expression after day 5 of the NPC time-course, is enriched for GO terms such as Notch signaling pathway (p < 1×10^−3^) and includes TFs like *Sox9* (Figure 2B). Cluster RR9, which increases in expression after day 3 of the DE time-course, is enriched for GO terms such as digestive tract morphogenesis (p < 2×10^−9^) and includes TFs like *Pitx2, Gata4, Sox17,* and *Gata6* (Figure 2B). We also performed a GO analysis of the five tan clusters of ESC high genes (Figure S3A). Cluster RR7, which decreases in NPC after day 2 and DE after day 3, is enriched for GO terms such as in utero embryonic development (p < 1×10^−5^). Cluster RR17, which decreases after day 4 in NPC and after day 3 in DE, is enriched for GO terms including blastocyst formation (p < 5×10^−4^). GO analysis of differentially expressed genes clustered by expression profile reveals that genes with shared functions are co-expressed during NPC and DE differentiation along with key TFs.

Next, we investigated changes in chromatin accessibility in the differentiation time-courses using ATAC-seq by clustering differential cCREs as previously described for gene expression (Figure 2C). For genes with more than one associated cCRE, each cCRE is analyzed independently, therefore a gene can have cCREs that are in different clusters. We detected 51,417 accessible chromatin regions in two or more replicates across both NPC and DE differentiation (Table S1). Of these, 25,665 differential cCREs grouped into 24 clusters of distinct patterns across time (Table S3). We observe a decrease in accessibility during differentiation in 9,375 (37%) of cCRE comprising nine clusters (labeled in tan; RA16, RA5, RA13, RA24, RA15, RA9, RA4, RA8, RA6) (Figure 2C). Clusters that increased in accessibility in the NPC time-course (labeled in blue; RA7, RA11, RA20, RA1, RA21, RA22) accounted for 5,124 (20%) of differential cCRE. Clusters that increased in accessibility in the DE time-course (labeled in green; RA10, RA14, RA2, RA3, RA19, RA12) accounted for 8,383 (33%) of cCRE. Clusters with shared changes in both time-courses (black; RA17, RA18, RA23) accounted for 2,762 of genes (10%) of cRE. Similar to the gene expression analysis, we calculated the correlation of the cCRE profiles over each differentiation time-course (Figure S2C) as well as plotted the samples on UMAP (Figure S2E). We found that UMAP1 represents time and UMAP2 represents the difference between NPC and DE differentiation. Both the correlation plot and UMAP suggest there are substantial changes in accessibility during differentiation (Figure S2B).

We then focused on the accessible regions surrounding TF genes that are also differentially expressed (Figure 2C). TFs found to be differential in both expression and accessibility profiles may be key in the regulation of differentiation for their respective cell types. Concordant cCREs have an accessibility profile that matches their gene expression profile in the specific cell-type. Discordant cCREs have an accessibility profile that does not match the gene expression profile. Examples of TFs found with concordant differential expression and chromatin accessibility include *Mef2a*, which increases in expression in NPC (RR8) and has one concordant cCRE (RA18) (Fig 2C). *Mef2a* has been shown to regulate neuron differentiation (Shalizi et al. 2006). Another TF that shows an increase in accessibility in NPCs is *bhlhe40* (RR2) (Figure 2C). There are three differential cCRE surrounding *bhele40*, which is needed for proper neuronal function (Hamilton et al. 2018). Each *bhele40* cCRE has a different accessibility profile, one of which is concordant (RA7) (Figure 2C). A TF that shows an increase in accessibility in DE is *Eomes* (RR9), which is essential for DE specification (Figure 2C) (Teo et al. 2011). There are five differential cCREs surrounding *Eomes*, four of which are concordant (RA2, RA14, RA19) (Figure 2C). Of the differentially expressed TFs, 63% have at least one cCRE that is concordant to its expression profile. This suggests that cell-type specific TF expression in rat is accompanied by differential accessibility of cCREs in a concordant direction, which we analyze in greater detail below.

To explore functional roles of cell-type specific TF cCREs we used GO for clusters increasing in NPCs (blue) and the DE (green) (Figure 2D). Cluster RA7, which increases in accessibility after day 1 in NPC differentiation, is enriched for GO terms like brain development (p < 2×10^−9^) and includes regions around important TFs like *Gbx2, Id1, Notch1, Rara,* and *Sox2* (Figure 2D). Cluster RA11, which increases in accessibility after day 2 of NPC differentiation, is enriched for GO terms such as nervous system development (p < 4×10^−22^). Cluster RA17, which increases in accessibility after day 2 in NPC differentiation, is enriched for GO terms including brain development (p < 4×10^−14^). Cluster RA10, which increases in accessibility after day 2 in DE differentiation, is enriched for GO terms such as urogenital system development (p < 5×10^−11^) and includes regions associated with *Rara* and *Foxb1*. Cluster RA19, which is highest in accessibility at day 4 of DE differentiation, is enriched for GO terms including tube formation (p < 1×10^−11^) and includes regions near *Gata3*. We also analyzed GO terms from each ESC-like cCRE cluster (Figure S3B). RA4, which gradually decreases until day 8 in NPC differentiation and after day 3 in DE differentiation, is enriched for GO terms including cell morphogenesis involved in differentiation (p < 5×10^−11^). Cluster RA13, which decreases after day 1 in NPC differentiation, is enriched for GO terms such as stem cell population maintenance (p < 1×10^−5^) and stem cell differentiation (p < 2×10^−6^).

We investigated which TF motifs are found in differentially accessible regions in each cluster using Homer (Figure S3C, S3D) (see methods). Motifs found in ESC-like clusters include Sox2, Pou5f1 (Oct4), and Nanog. DE cluster motif enrichments include; Gata4, Gsc, Tead2, Foxa1, Eomes, and Sox17 (Figure S3C). Many motifs were enriched in both cell-types. The Gsc motif is enriched in two clusters (RA2, RA23 both p < 1×10^−25^). The Eomes motif was enriched in eight clusters (RA1, RA2, RA14, RA15, RA17, RA19, RA21, RA24 all p < 5×10^−19^), three of these clusters are in the DE group. NPC cluster motif enrichments include Tead4, Sox2, and Maz. As expected, we find the Sox2 motif in nine clusters (RA1, RA4, RA6, RA7, RA8, RA9, RA11, RA13, RA15 all p < 5×10^−26^), including the ESC and NPC groups, suggesting Sox2 is required for regulation of both ESC maintenance and NPC differentiation (Shuchen Zhang and Cui 2014). The Tcf12 motif was enriched in 3 clusters (RA9, RA19, RA21 all p < 2×10^−18^) and has been shown to be important in NPC differentiation (M. Uittenbogaard 2002; Simone Mesman 2017). In conclusion, GO term analysis paired with *de novo* motif enrichment highlights TFs with binding sites in cell-type and time-course specific cCREs.

To mine clusters for differential gene expression and cCRE interactions we analyzed the relationship between differential RNA and ATAC clusters in rat. We used the overlaps to build a contingency table that contains the association of each of the differentially accessible regions with the mRNA clusters (see methods, Table S4). We detected 55 sets of significantly enriched interactions between ATAC and RNA clusters. Of these, 22 have concordant significantly enriched interactions with both the expression and accessibility higher in the same cell-type (NPC (blue), DE (green), or ESC (tan)). 29 are discordant significantly enriched interactions (i.e. expression is high in NPC and accessibility is high in DE), and an additional four significant interactions in which the accessibility is not associated with any cell-type (black; Cluster RA23). We focused on cCREs surrounding TFs in each significantly overlapping cluster and found 231 instances of cCREs for TFs (totaling 109 TFs). Some of the TFs that have a shared affiliation with NPC clusters include Mef2a (RR8, RA18) and bhlhe40 (RR2, RA7). Some of the TFs with shared DE affiliations include T (RR9, RA2), Foxa2 (RR9, RA2, RA9, RA19), Sox17 (RR9, RA14), and Foxb1 (RR5, RA3, RA10, RA14). TFs in ESC affiliated clusters are Tcf7l1 (RR12, RA15), Nanog (RR12, RA9, RA15), and Foxp1 (RR7, RA15). All of the TFs listed above have increased expression and promoter accessibility in their respective cell types and probably play an important role in the regulation of differentiation in rat. We investigated the TFs that have significant overlapping clusters from different groups (i.e. increased expression in NPC and increased accessibility in DE) between DE and NPC clusters such as Tgif1 (RR5, RA17), Notch1 (RR5, RA11,17,18), Id1 (RR5, RA18), and Jun (RR5, RA11). Some of these genes have important roles in the regulation of differentiation into one or both cell-types in human (David Wotton 2018; Pierfelice et al. 2008; Chu et al. 2016; Boareto, Iber, and Taylor 2017), but have not been previously shown in rat. These TFs either have increased expression, or increased promoter accessibility, in one of the time-courses suggesting that these TFs are candidate regulatory genes for at least one of the differentiation time-courses in rat.

We then mined the accessible regions with significant overlap (from Figure 2E) for TF footprints (see methods, Figure 2F). The most enriched footprints are Ctcf, Sp5, Klf, Sp1, and Zic, which are found in multiple ESC, DE, and NPC sets of enriched interactions and are probably needed for the regulation of differentiation of both lineages as well as early differentiation and ESC maintenance. Footprint enrichments for Atf3 are mostly exclusive to significant NPC overlaps, whereas Sox17 and Eomes footprints are mainly exclusive to significant DE cluster overlaps. As expected, Nanog and Sox2 have significant footprint enrichment in ESC cluster overlaps. Jund and Junb footprints are found in clusters with significant overlapping clusters with NPC and DE affiliations. Although we expected a higher fraction of concordant significant overlapping clusters with matching gene expression and chromatin accessibility for their respective cell types, we still have many candidate regulatory genes for each cell type.

### Distinct Transcriptional and Chromatin Accessibility Trajectories for NPC and DE in Human

We repeated the same analysis for human ES cell differentiation using the human H1 ES cell line to identify candidate regulators for the differentiation of each cell type. We collected 27 RNA-seq human datasets (see methods), resulting in 17,661 genes expressed over 1 TPM in two or more replicates across both NPC and DE time-courses (Table S1). We used maSigPro to identify 5,755 differentially expressed genes grouped into 18 distinct clusters (Figure 3A, Table S5). Four clusters containing 1,338 genes (23%) decrease over time (HR8, HA13, HA9, HA2). Seven clusters comprising 1,638 genes (29%) increase during NPC differentiation (HR15, HA11, HA16, HA14, HA4, HA12, HA18). Another seven clusters containing 2,779 genes (48%) increase during DE differentiation (HR10, HA7, HA1, HA17, HA6, HA5, HA3). We plotted the correlation of the expression profiles over each differentiation time-course (Figure S4A) as well as plotted the samples on a UMAP (Figure S4B). UMAP1 represents the difference between the two differentiation time-courses, and UMAP2 represents differentiation time with larger changes in the differentially expressed genes later in each time-course suggesting a clear difference between early and late differentiation as seen in rat (Figure S4A-C). Clusters that increased during NPC differentiation included TFs important in neurodevelopment and expression in early neurons such as *SOX2* (HR11), *FOXP1* (HR15), *PAX6* (HR16), *MEIS2* (HR14), *MEF2A* (HR16), and *JUN* (HR12) (David Wotton 2018; Moore et al. 2013)(Ziller et al. 2015)(David Wotton 2018; Moore et al. 2013). Clusters that increased during DE differentiation included TFs that are important in endoderm development and early differentiation including *SOX17* (HR5), *GATA6* (HR5), *FOXA2* (HR3), *GATA4* (HR3), *GSC* (HR6), *EOMES* (HR17) (Woodland and Zorn 2008; Teo et al. 2011; Sinner et al. 2004; Shimosato, Shiki, and Niwa 2007). The ESC-like clusters, which decrease in expression in both of the time-courses, include interesting TFs involved in ESC maintenance including *NANOG* (HR2), *OCT4* (HR2), and *TCF7L1* (HR2). Many of these TFs are shared between rat and human; some examples are, *PAX6, JUN, MEF2A,* and *MEIS2* for NPC differentiation and *GATA4, GATA6, LHX1,* and *SOX17* for DE differentiation. The TFs found in cell-type specific clusters in human suggest they are expressed in the appropriate pattern to regulate differentiation into either NPC or DE.

**Figure 3:**
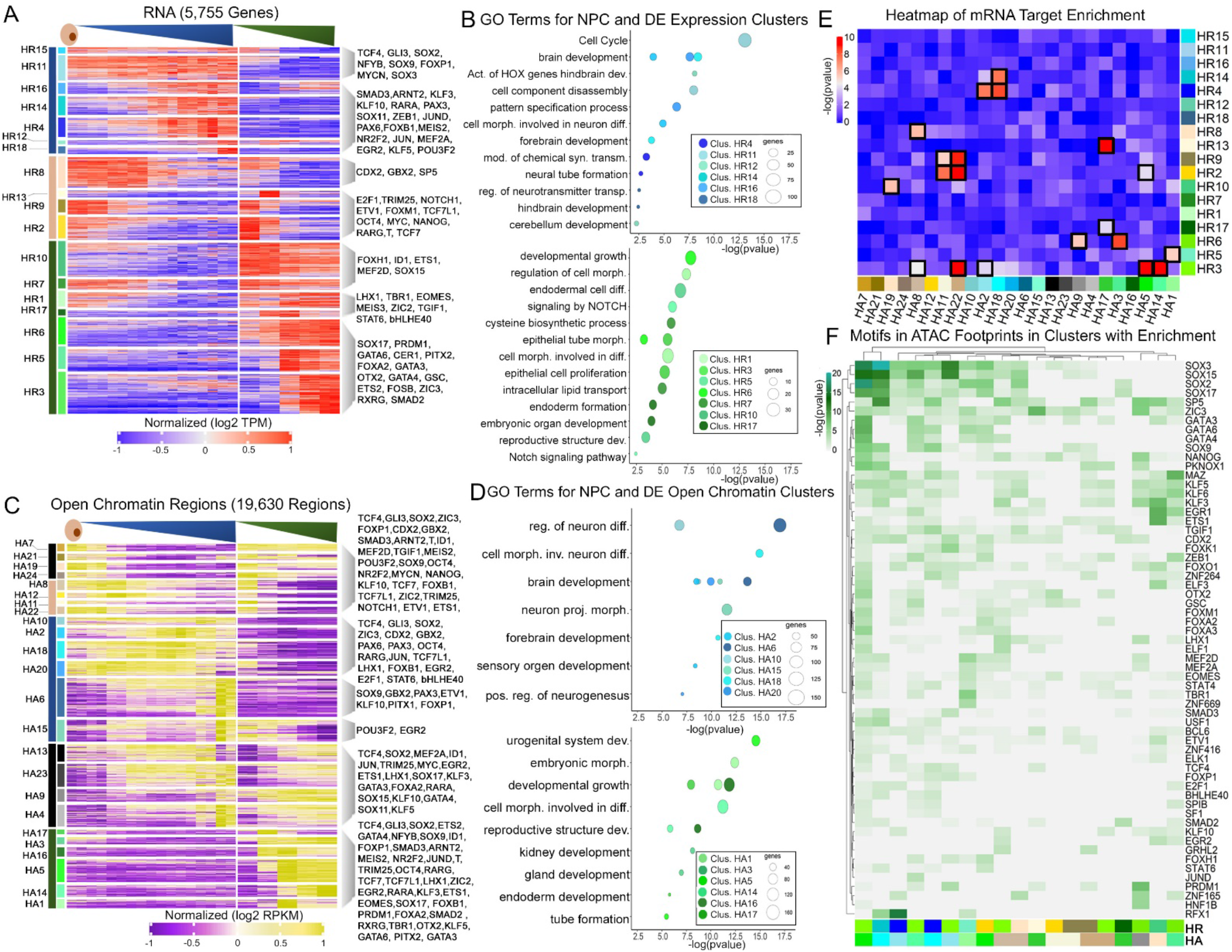
Distinct Transcriptional and Chromatin Accessibility Trajectories for NPC and DE in Human. A) Heatmap of 5,755 differentially expressed genes (alpha = 0.05, p < 0.05) in NPC and DE differentiation in human Hierarchical clustering was used to generate eighteen expression clusters denoted by the adjacent number. Transcripts per million (TPM) values were log transformed and row-mean normalized for all genes. Colors on the far left correspond to the group (NPC, DE, or ESC) the cluster is associated with. The second color bar is for each individual cluster. Select TFs are labeled. B) GO terms for NPC and DE related gene expression clusters. The GO terms for clusters that increase over the NPC time course are in blue. The GO terms for clusters that increase over the DE time course are in green. GO terms are determined using metascape. C) Heatmap of 19,630 differential cCRE (alpha = 0.05, FDR = 0.05, p < 0.05) in NPC and DE differentiation in human. Hierarchical clustering was used to generate twenty four DNA region clusters denoted by the adjacent number. RPKM (Reads Per Kilobase Million) values were log transformed and row-mean normalized for all regions. D) GO terms for NPC and DE related cCRE clusters. The GO terms for clusters that increase over the NPC time-course are in blue. The GO terms for clusters that increase over the DE time-course are in green. GO terms are determined using metascape. E) Heatmap of X^2^ p-values on a contingency table of clusters of accessible regions and expression clusters with applied P-value of 10^−4^ (Bonferroni corrected P-value: 0.05/[18 × 24]). The significant cluster overlaps are surrounded by a black box. The colors next to each cluster name match the colors in Figures 3A and 3C. F) ATAC footprints in regions of significant overlap between clusters. The color bars are the same in A, C, and E for each RNA and ATAC cluster.

We performed GO analysis of clusters increasing in NPC (blue) or DE (green) (Figure 3B). Cluster HR12, whose genes increase in expression after day 7 in NPC differentiation, is enriched for GO terms like cerebellum development (p < 5×10^−3^) and includes *RARA*. Cluster HR14, which increases in expression after day 3 in the NPC time-course, is enriched for GO terms such as brain development (p < 4×10^−9^) and forebrain development (p < 1×10^−4^) and includes *MEIS2*. Cluster HR5, which sharply increases in expression on day 3 followed by a gradual decrease during DE differentiation, is enriched for GO terms such as endodermal cell differentiation (p < 1×10^−7^) and includes *GATA6* and *SOX17*. Cluster HR17, which has high expression on days 2 and 3 of the DE time-course, is enriched for GO terms such as endoderm formation (p < 7×10^−5^) and includes *EOMES* and *LHX1*. Next, we considered clusters in the ESC group (tan, Figure S5A). For example, Cluster HR2, which decreases after day 1 in both NPC and DE differentiation, is enriched for GO terms such as regulation of growth (p < 7×10^−7^). Cluster HR13, which decreases after day 1 in NPC differentiation and is highest in day 2 in DE differentiation, is enriched for GO terms such as embryonic morphogenesis (p < 4×10^−6^).

Next, we investigated changes in chromatin accessibility in the human differentiation time-courses using ATAC-seq (Figure 3C). We collected 27 ATAC-seq human datasets (see methods), resulting in 75,271 accessible chromatin regions in two or more replicates (Table S1). We clustered temporally differential cCRE as performed in rat. We found 19,630 differentially accessible cCRE that grouped into 24 clusters (Figure 2C, Table S6). We observed a loss in accessibility in 4,636 (24%) of cCRE that form eight clusters (HA7, HA21, HA19, HA24, HA8, HA12, HA11, HA22). We observed six clusters that increased in chromatin accessibility in the NPC lineage (labeled in blue; HA10, HA2, HA18, HA20, HA6, HA15) accounting for 4,828 (25%) of differential cCRE. We found six clusters that increased in accessibility in the DE time-course (HA17, HA3, HA16, HA5, HA14, HA1), totaling 8,480 (42%) of the differential cCRE. Four clusters shared changes in accessibility in the two time-courses (black; HA13, HA23, HA9, HA4) totaling 1,686 (9%) of the differential cCRE. We plotted the correlation of cCRE profiles for each cell type (Figure S4D) and plotted the differential cCRE from each sample on a UMAP (Figure S4E). UMAP1 represents differentiation time and UMAP2 represents the difference between cell types. Similar to what we found in rat, human differential cCRE show substantial differences between early and late stages (Figure S4B). We then focused on cCREs near TFs that are also differentially expressed to find which cCREs show increasing or decreasing accessibility and expression in the same direction (concordant) or different directions (discordant) changes in both techniques (Figure 3C). TCF4, a TF that supports differentiation into neurons, expression increases (HR11) during human NPC differentiation (Mesman, Bakker, and Smidt 2020). TCF4 has 3 out of 8 associated differential cCREs (HA16, HA5, HA17, HA11, HA20, HA18) and 3 of these cCREs (HA18, HA20) with concordant increases in accessibility to the RNA expression profile. *GBX2*, a neuroectodermal marker, increases in expression over the NPC time-course and has 4 out of 9 concordant differential cCRE (HA6, HA18, HA20) (Martinez-Barbera et al. 2001). *RXRG*, which is involved in endoderm development (Sherwood, Chen, and Melton 2009), increases in expression over the DE time-course (HR6) and has two differential cCREs, one of which concordant (HA1). Of the differential TF cCREs, 66% have at least one cCRE that is concordantly changing with its expression profile. Similar to what we found in rat, this suggests that TFs found in cell-type specific expression clusters show concordant differential accessibility of at least one associated cCRE.

Next, to explore functional roles of cell-type specific TF cCREs we analyzed each cCRE cluster using GO for clusters increasing in NPC (blue) and DE (green)(Figure 3D). Cluster HA2, which has the greatest accessibility on days 5 and 6 in NPC differentiation, is enriched for GO terms like brain development (p < 3×10^−9^) and includes *E2F1, LHX1, PAX6,* and *RARA*. Cluster HA6, which gradually increases from days 2 to 8 in NPC differentiation, is enriched for GO terms such as regulation of neuron differentiation (p < 1×10^−17^) and includes *LEF1*, and *SOX9*. Cluster HA18, with the greatest accessibility on day 5 of the NPC time-course, is enriched for GO terms such as forebrain development (p < 2×10^−11^) which includes *GBX2, SOX2, LEF1,* and *FOXB1*. Cluster HA3, which increases in accessibility after day 1 of DE differentiation, is enriched for GO terms like gland development (p < 1×10^−7^) and includes *PITX2*, *RARA* and *RARG*. Cluster HA14, which increases after day 3 in DE differentiation, is enriched for GO terms like reproductive structure development (p < 1×10^−6^) and includes *GATA4* and *RARA*. Many of the TFs upregulated in both cell-types are upregulated in both rat and human, such as *LHX1, bHLHE40, SOX9, SOX2*, and *RARA* for NPC differentiation and, *GATA3, GATA4, PITX2, RARA, CER1,* and *SOX17* for DE differentiation (Figure 2D and 3D) and are important for regulation of differentiation into their specific cell-types (Dollé 2009; Symmank et al. 2019; Scott et al. 2010; Hamilton et al. 2018; Shimosato, Shiki, and Niwa 2007; Faucourt et al. 2001). We also used GO to investigate ESC specific clusters (Figure S5B). Cluster HA11, which decreases in accessibility after day 1 in both NPC and DE differentiation, is enriched for GO terms like cell morphogenesis involved in differentiation (p < 3×10^−7^) and stem cell population maintenance (p < 9×10^−6^). Cluster HA12, which decreases in expression after day 3 in NPC differentiation and day 1 in DE differentiation, is enriched for GO terms like embryonic morphogenesis (p < 8×10^−9^). GO analysis of cCRE clusters shows that sets of TFs with shared functions are increasing in accessibility in similar temporal patterns in human and many of these regions show similar patterns in rat.

We then investigated which *de novo* motifs are found in each cluster using Homer (Figure S5C-D). We found POU5F1, SOX2, NANOG, and FOXP1 motif enrichment in ESC clusters, whereas NPC clusters were enriched for MEIS1, MEIS2, MEF2A, KLF, MAZ, RUNX, and SP1 motifs. The MEF2A motif is enriched in two clusters (HA2, HA20 both p < 1×10^−38^), both are NPC clusters. Motifs enriched in DE clusters include EOMES, NANOG, GATA4, OTX2, GSC, SOX17, and TEAD4 (Figure S5C). In conclusion, *de novo* motif enrichment highlights TFs with binding sites found in differential cCRE with time-course specific functionality in human.

Next, we analyzed the relationship between differential cCREs and expression clusters in human, similar to what we did in rat (see methods, Figure 3E). We detected 20 sets of significantly enriched interactions (Table S7). Of these, 13 had concordant significantly enriched clusters, with expression and accessibility clusters falling into the same category, NPC (blue), DE (green), or ESC (tan). Seven discordant significantly enriched clusters, and one significant interaction in which the accessibility cluster is unaffiliated with a specific cell-type (black). We focused on cCREs surrounding TFs in each significantly overlapping clusters and found 93 instances of cCREs for TFs (totaling 50 unique TFs). Examples of TFs with shared affiliations with NPC clusters include *FOXB1* (HR4, HA18) and *PAX8* (HR4, HA18). Interestingly, these two TFs are in the same RNA cluster and have cCREs that are in the same ATAC clusters, which suggests coordinated regulation in NPC differentiation. Some of the TFs with shared DE affiliations include *GATA4* (HR3, HA5), *SOX17* (HR5, HA1), *PITX2* (HR6, HA3), *FOXA2* (HR3, HA5), and *OTX2* (HR3, HA5). Of note, many of the cCREs with significant overlaps in the DE group had multiple cCRE regions with the same cluster overlap identity, *GATA4, SOX17, PITX2,* and *OTX2* all had multiple regions in significant overlapping clusters with the same cCRE cluster affiliations. TFs in ESC affiliated clusters include *POU5F1* (HR2, HA22). All of the TFs listed above have increased expression and promoter accessibility in their respective cell types, making them clear candidates for regulatory activity in their respective differentiations. We investigated the TFs that have significant overlapping clusters from different groups (i.e. increased expression in NPC and increased accessibility in DE) between DE and NPC clusters such as *RXRG* (HR3, HA2). This TF has increased expression in the DE time-course and increased accessibility in the NPC time-course in human, suggesting it is a strong regulatory factor candidate for at least one of the differentiation time-courses.

We then mined the ATAC regions with significant overlaps from Figure 3E for footprints (see methods, Figure 3F). The most enriched footprints include SOX2, SOX17, and SP5. These motifs are found in multiple ESC, DE, and NPC enriched interactions and are probably needed for the regulation of differentiation of both lineages and ESC maintenance. Footprint enrichments for E2F1 and bHLHE40 are mainly exclusive to significant NPC cluster overlaps. EOMES, OTX2, GATA3, and GSC that are mainly exclusive to significant DE cluster overlaps. NANOG has motifs in ESC cluster overlaps. Whereas SMAD3 and FOXP1 motifs are found in clusters with significant overlapping clusters with NPC and DE affiliations. Since the majority of the significantly overlapping clusters are concordant, we have many cCREs for TFs for each cell type. There are a few discordant cCREs, which are also interesting to investigate because they have differing promoter accessibility and expression in their respective cell types. The discordant cCREs may still be involved in the regulation of differentiation possibly via repression.

### Distinct Transcriptional and Chromatin Accessibility Trajectories for NPC Between Rat and Human

We then focused on one-to-one orthologs that are expressed in at least one species and detected 9,898 genes expressed above 5 TPM in at least two replicates across NPC differentiation in either species. We computed the differential expression between rat and human during differentiation into NPC using maSigPro (Figure 4A). We found 3,562 differentially expressed orthologs grouped into 18 clusters (Table S8). We separated the differentially expressed clusters by early, late, and species-specific expression profiles. For the early group, expression profiles were highest in both human and rat during the first half of the time-courses (roughly from days 0 to 5). We found four clusters accounting for 1,244 genes (35%) that had high expression in the early group (light blue; JNR10, JNR1, JNR7, JNR18, JNR14). For the late group, both human and rat expression profiles peaked in expression later in the NPC time-course (from days 4 to 8). We found six clusters accounting for 1,117 genes (31%) that had increased in expression in the late group (dark blue; JNR5, JNR16, JNR8, JNR11, JNR3, JNR17). For the species-specific group, expression profiles had an opposite trend in each species (i.e. early group in rat, and late group in human). We determined seven clusters totaling 1,201 genes (34%) have species-specific expression profiles (black; JNR2, JNR6, JNR4, JNR9, JNR13, JNR15, JNR12). Two of these clusters have high expression in the late group in rat and the early group in human (JNR2, JNR6). The other five clusters are in the early group in rat and the late group in human (JNR4, JNR9, JNR13, JNR15, JNR12). We focused on TFs found in each cluster as they may play a role in the regulation of NPC differentiation in both species. TFs found in the early group include important ESC maintenance genes like *NANOG* (JNR14), *POU5F1* (JNR10), and NRSF/*REST* (JNR14), and important early neuronal differentiation genes, *ZIC3* (JNR1), *TGIF1* (JNR10), and *GBX2* (JNR14) (Martinez-Barbera et al. 2001; David Wotton 2018). TFs found in the later expression clusters like, *JUND* (JNR5), *SOX11* (JNR5), *MEIS3* (JNR8), *JUN* (JNR8), *JUNB* (JNR8), *ID1* (JNR8), *RARG* (JNR3), *RARA* (JNR5), *TEAD2* (JNR3), *PBX1* (JNR3), and *SALL2* (JNR3), are more likely to be involved in the final stages of NPC differentiation or NPC maintenance (Golonzhka et al. 2015). The TFs in the species-specific clusters are interesting because they might have differing regulation in each species. These TFs include; *SOX2* (JNR4), *LEF1* (JNR12), *CTCF* (JNR15), *TCF4* (JNR15), *MEIS2* (JNR13), *KLF2* (JNR9), *KLF3* (JNR13), *SP1* (JNR15), and *ID3* (JNR6) (Okamoto et al. 2002). While 66% of expressed orthologs show conserved changes in expression between the two species, another 34% show species-specific behavior. The TFs found in each of the three groups are expressed in the appropriate temporal pattern to regulate different temporal stages of NPC differentiation in one or both species.

**Figure 4:**
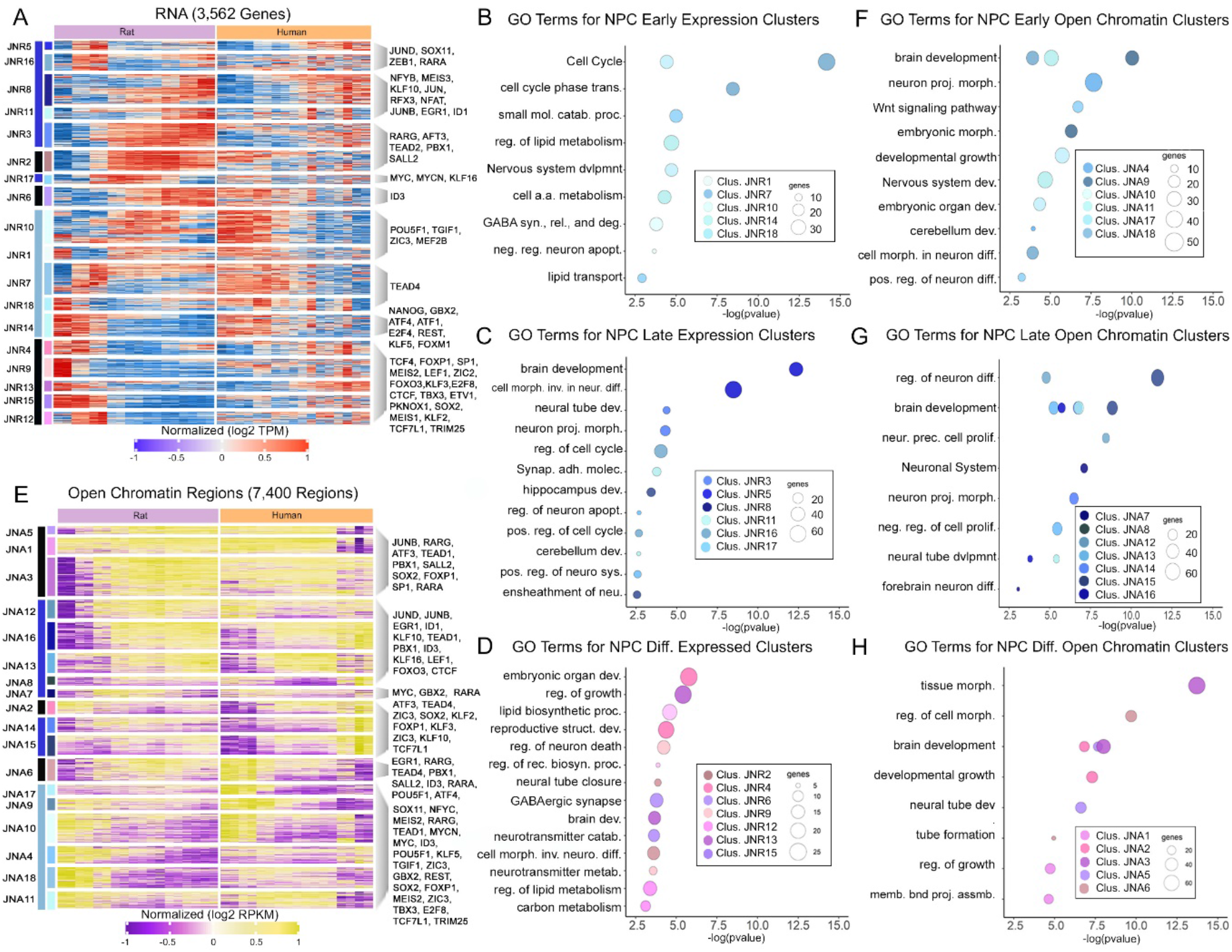
Distinct Transcriptional and Chromatin Accessibility Trajectories for NPC Between Rat and Human. A) Heatmap of 3,562 differentially expressed genes (alpha = 0.05, p < 0.05) in NPC differentiation in rat and human. Hierarchical clustering was used to generate eighteen expression clusters denoted by the adjacent number. Transcripts per million (TPM) values were log transformed and row-mean normalized for all genes. Colors on the far left correspond to the group (early [light blue], late [dark blue], or species-specific [black]) the cluster is associated with. The second color bar is for each individual cluster. Select TFs are labeled. B) GO terms for Early grouped gene expression clusters. The GO terms for clusters of genes that increase early in both rat and human NPC differentiation are in light blue. GO terms are determined using metascape. C) GO terms for Late grouped gene expression clusters. The GO terms from genes that increase later in the NPC time-course for both rat and human are in dark blue D) GO terms for species-specific grouped gene expression clusters. These genes showed different expression profiles in rat and human. GO terms were obtained from genes that differ during differentiation of the two species. E) Heatmap of 7,400 differential cCRE (alpha = 0.05, FDR = 0.05, p < 0.05) in NPC differentiation in rat and human. Hierarchical clustering was used to generate 18 DNA region clusters denoted by the adjacent number. RPKM (Reads Per Kilobase Million) values were log transformed and row-mean normalized for all regions. F) GO terms for cCRE that were accessible early on in NPC differentiation in both rat and human. G) GO terms for cCRE that were accessible later on in NPC differentiation for both rat and human. H) GO terms for cCRE that have species-specific accessibility in NPC differentiation between rat and human.

Next, we investigated each cluster using GO and separated clusters by their group affiliation, starting with the early group (Figure 4B). Early cluster JNR1 is enriched for GO terms such as negative regulation of neuron apoptotic process (p < 2×10^−4^) and GABA synthesis, release, reuptake, and degradation (p < 1×10^−4^). Early cluster JNR10 is enriched for GO terms such as nervous system development (p < 1×10^−5^). We next investigated GO terms for clusters from the late group (Figure 4C). Late cluster JNR3 is enriched for GO terms such as neural tube development (p < 4×10^−5^) and includes *RARG, SALL2,* and *TEAD2*. Late cluster JNR8 is enriched for GO terms such as cell morphogenesis involved in neuron differentiation (p < 3×10^−9^) which includes *JUN*. Next we investigated GO terms found for the clusters found in the species-specific group (Figure 4D). Species-specific cluster JNR9, which is very highly expressed in day 0 in rat, with a sharp decrease on day 1 and increases after day 4 in human, is enriched for GO terms such as regulation of neuron death (p < 5×10^−5^). Species-specific cluster JNR15, which starts with high expression and decreases after day 2 in rat, and increases after day 3 in human, is enriched for GO terms such as neurotransmitter catabolic process (p < 2×10^−4^). GO analysis of gene expression indicates species-specific clusters with differential expression between the two species during NPC differentiation are enriched for a distinct subset of neuronal GO terms.

Next, we compared the changes in chromatin accessibility in NPC differentiation between rat and human (Figure 4E) by clustering cCREs as previously described. We detected 25,599 uniquely alignable accessible chromatin regions in at least two replicates across NPC differentiation in each species. We found 7,400 differential cCRE that grouped into 18 clusters (Table S9). We classified the cCRE clusters into three different groups: early, late, and species-specific. We found six clusters accounting for 2,444 cCRE (33%) that were the most accessible in the early group (JNA17, JNA9, JNA10, JNA4, JNA18, JNA11). There are seven clusters accounting for 2,708 cCRE (37%) in the late group (JNA12, JNA16, JNA13, JNA8, JNA7, JNA14, JNA15). There are five clusters consisting of 2,248 cCRE (30%) in the species-specific group (JNA5, JNA1, JNA3, JNA2, JNA6). Clusters in the early group included cCREs surrounding TFs that are a mix of ESC markers, including *POU5F1* (JNA6, JNA9), *TCF7L1* (JNA2, JNA9), and NRSF/*REST* (JNA18), and early neuronal differentiation markers, including; *MEIS2* (JNA9, JNA18), *MYCN* (JNA9, JNA10), *TGIF1* (JNA9, JNA10, JNA11, JNA18), *FOXP1* (JNA1, JNA9, JNA10, JNA14), and *ZIC3* (JNA2, JNA9, JNA11, JNA18). cCREs in the late group include TFs that are important in the final stages of NPC differentiation in both species; including, *JUND* (JNA13), *JUNB* (JNA3, JNA13), *EGR1* (JNA6, JNA13), *ID3* (JNA6, JNA10, JNA12, JNA16), *GBX2* (JNA7, JNA9), *TEAD4* (JNA6, JNA15), *ID1* (JNA12, JNA13), and *LEF1* (JNA13). Arguably, the most interesting are cCREs in the species-specific group, as these cCREs may be important in species-specific aspects of differentiation in each species. The cCREs in species-specific clusters include *RARA* (JNA3, JNA6, JNA7), *RARG* (JNA3, JNA6, JNA10), *SOX2* (JNA2, JNA3, JNA9), *PBX1* (JNA3, JNA6, JNA12), *ATF3* (JNA3, JNA15), *KLF3* (JNA2), and *SALL2* (JNA5, JNA6). While 70% of alignable cCRE show conserved changes in chromatin accessibility between the two species, another 30% show species-specific behavior.

Next, we used GO to analyze each cCRE cluster, separating the clusters into early (light blue), late (dark blue) and species-specific (pink), starting with the early group (Figure 4F). Early cluster JNA4 is enriched for GO terms such as neuron projection morphology (p < 2×10^−8^) which includes cCREs for MEF2C. Early cluster JNA9 is enriched for GO terms such as brain development (p < 9×10^−11^) and embryonic morphology (p < 5×10^−7^) which include cCREs for *GBX2, MEIS2, OTX2, SOX2, POU5F1, ZIC3, TGIF2,* and *FOXP1*. Clusters in the late group were analyzed for GO terms (Figure 4G). Late cluster JNA 16 is enriched for GO terms such as regulation of neuron differentiation (p < 2×10^−12^) and includes cCREs for *SOX11, NOTCH1,* and *MEF2A*. Late cluster JNA13 is enriched for GO terms such as neural precursor cell proliferation (p < 3×10^−9^) and includes cCREs for *LEF1* and *SOX5*. These cCREs are accessible in a pattern conducive to conserved regulation in NPC differentiation in rat and human. We then focused on species-specific clusters (Figure 4H). Species-specific cluster JNA3, which increases in accessibility in rat after day 1 and decreases in human after day 6, is enriched for GO terms such as brain development (p < 2×10^−8^) and tissue morphogenesis (p < 2×10^−14^) and includes *LHX1, NOTCH1, RARA, SOX2, GATA4, PBX1,* and *RARG*. Cluster JNA5, which increases in accessibility in rat after day 0 and decreases in human after day 6, is enriched for GO terms such as brain development (p < 2×10^−8^) and neural tube development (p < 2×10^−7^) that includes cCREs such as SALL2. Since these cCREs differ in their temporal accessibility in each species, they may play species-specific roles during NPC differentiation, while cCREs in early or late clusters may play a more conserved role in the same process.

### Distinct Transcriptional and Chromatin Accessibility Trajectories for DE Between Rat and Human

We analyzed the expression one-to-one orthologs during DE differentiation in both species using maSigPro and detected 7,320 differentially expressed genes (Figure 5A). These genes were further classified into 12 clusters with distinct stem, mesendodermal, or definitive endodermal profiles (Figure 5A, Table S10). We separated the clusters by early, late, and species-specific expression profiles. We found 2,691 genes (37%) in five clusters (JDR1, JDR2, JDR8, JDR9, JDR10) that had increased expression in human and rat DE time-courses early in DE differentiation (light green). JDR1, JDR2, and JDR9 showed similar expressions in both human and rat, whereas JDR8 and JDR10 showed stronger expression in human. These clusters were enriched with pluripotent markers such as *POU5F1* (JDR1), *NANOG* (JDR9) and *SOX2* (JDR9) in both human and rat (Figure 5A). We found 2,666 (36%) genes in four clusters (JDR3, JDR6, JDR7, JDR11) with increased later expression (dark green). JDR3, JDR7, and JDR11 showed similar expression in both species whereas JDR6 is more strongly expressed in rat. Clusters with genes expressed at the late stage were highly enriched with endodermal markers that showed gradual activation towards the end of DE differentiation. TFs found in the late group included *GATA6* (JDR7), *EOMES* (JDR11), *FOXA2* (JDR11), *GATA4* (JDR11), *SOX17* (JDR11), and *GSC* (JDR11). Brachyury/T (JDR6), which is a mesendodermal marker (Faial et al. 2015; Turner et al. 2014), had the highest expression level at day 2 differentiation for human and day 4 differentiation for rat and then maintained the expression level towards the end of time-course. Similar to what we observed in the NPC differentiation, we also identified species-specific expression group profiles between two species. The remaining 1963 (27%) genes in three clusters (JDR4, JDR5, JDR12) have more pronounced species-specific expression profiles (black). JDR5 and JDR12 exhibit early expression in rat and late expression in human while cluster JDR4 has late activation in rat and early activation in human. These clusters include a set of TFs associated with cell cycle regulation and biosynthetic processes were enriched in species-specific clusters such as *NRSF/REST* (JDR5), *ETS1* (JDR5), *ID3* (JDR4), *SALL2* (JDR4), and *RARG* (JDR4). While 73% of expressed orthologs show conserved changes in expression between the two species, another 27% show species-specific behavior.

**Figure 5:**
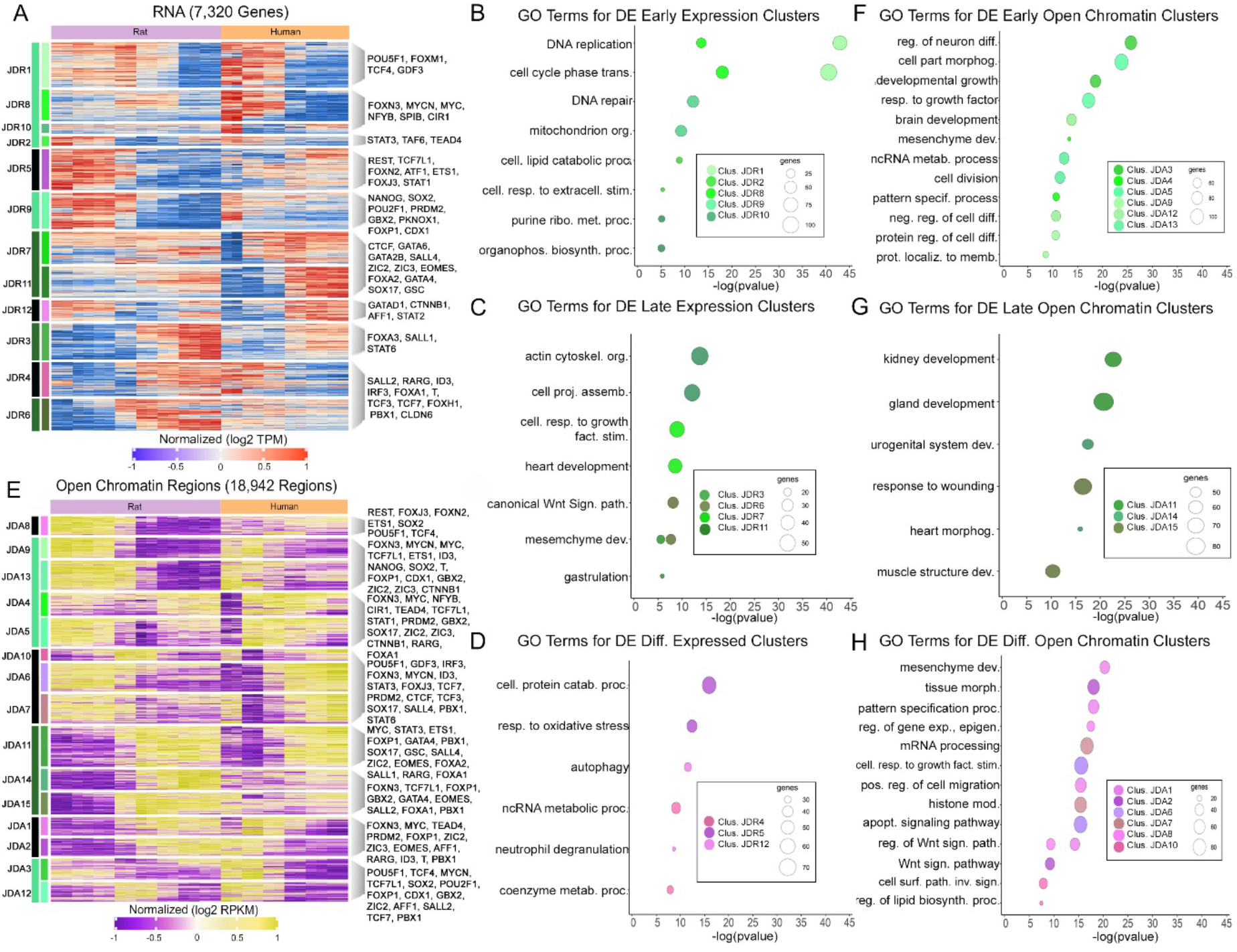
Distinct Transcriptional and Chromatin Accessibility Trajectories for DE Between Rat and Human. A) Heatmap of 7,320 differentially expressed genes (alpha = 0.05, p < 0.05) in DE differentiation in rat and human. Hierarchical clustering was used to generate 12 expression clusters denoted by the adjacent number. Transcripts per million (TPM) values were log transformed and row-mean normalized for all genes. Colors on the far left correspond to the group (early [light green], late [dark green], or species-specific [black]) the cluster is associated with. The second color bar is for each individual cluster. Select TFs are labeled. B) GO terms for Early grouped gene expression clusters. The GO terms for clusters of genes that increase early in both rat and human DE differentiation are in light green. GO terms are determined using metascape. C) GO terms for Late grouped gene expression clusters. The GO terms from genes that increase later in the DE time-course for both rat and human are in dark green. D) GO terms for species-specific grouped gene expression clusters. These genes showed different expression profiles in rat and human. GO terms were obtained from genes that differ during differentiation of the two species. E) Heatmap of 18,942 differential cCRE (alpha = 0.05, p < 0.05) in DE differentiation in rat and human. Hierarchical clustering was used to generate 15 DNA region clusters denoted by the adjacent number. RPKM (Reads Per Kilobase Million) values were log transformed and row-mean normalized for all regions. Colors on the far left correspond to the group (early [light green], late [dark green], or species-specific [black]) the cluster is associated with. The second color bar is for each individual cluster. Select TFs are labeled. F) GO terms for cCRE that were accessible early on in DE differentiation in both rat and human. The GO terms for clusters that increase over the DE time-course are in light green. G) GO terms for cCRE that were accessible later on in DE differentiation for both rat and human. The GO terms for these regions are shades of dark green. H) GO terms for cCRE that have species-specific accessibility in DE differentiation between rat and human. These regions are shades of purple and pink.

In order to determine the roles of differentially expressed genes during DE differentiation, we ran GO analysis on each cluster and summarized the biological processes of these genes by their early or late expression profiles. The early groupings of genes were significantly associated with DNA replication (JDR1, p < 1×10^−43^), cellular lipid catabolic process (p < 1×10^−8^)/response to the extracellular stimulus (p < 1×10^−5^) (JDR2) and DNA repair (JDR9, p < 1×10^−11^) in both human and rat (Figure 5B). Clusters JDR3,7,11 were highly enriched with endodermal markers that showed gradual activation towards the end of DE differentiation in both species. Genes in cluster JDR7 are associated with growth factor stimulus (p < 1×10^−8^) and heart development (p < 1×10^−8^) while genes in cluster JDR11 were significantly involved in gastrulation (p < 1×10^−5^) and mesenchyme development (p < 1×10^−5^) (Figure 5C). However, genes in cluster JDR3 were involved in basic cellular processes, such as actin cytoskeleton organization (p < 1×10^−13^) and cell projection assembly (p < 1×10^−12^) (Figure 5C). As mentioned above, we found genes in cluster JDR6 reached to the highest level during the intermediate stage of DE differentiation and Brachyury T marked this cluster as a mesendodermal cluster (Figure 5A). As expected, genes in JDR6 were significantly associated with canonical Wnt signaling pathway (p < 1×10^−8^), mesenchyme development (p < 1×10^−7^) and mesoderm development (p < 1×10^−7^) (Figure 5C). We also found three clusters that show reverse expression patterns in two species (Figure 5A). Genes in cluster JDR4 shown increased expression in rat but decreased in human were enriched in ncRNA metabolic process (p < 1×10^−9^) while genes in cluster JDR5 and JDR12 that were down-regulated during rat differentiation but up-regulated in human were highly involved in cellular protein catabolic process (p < 1×10^−15^), response to oxidative stress (p < 1×10^−12^), autophagy (p < 1×10^−11^) and neutrophil degranulation (p < 1×10^−8^) (Figure 5D).

To further understand transcriptional control during DE differentiation in human and rat, we identified cCREs that showed differentially chromatin accessibility during DE differentiation in two species (Figure 5E) and found 18,942 (74% out of 25,599) regions that formed 15 clusters with distinct accessibility patterns (Figure 5E, Table S11). We further selected clusters based on their associated genes that were shared in human and rat. Corresponding to the gene expression patterns (Figure 5A), we found that stem cell specific clusters (JDA3,9,13) enriched with pluripotent markers while clusters JDA11,14,15 were enriched with endodermal markers. This resulted in a loss of 3,856 (20%) regions while a gain of 4,506 (24%) regions during DE differentiation that were shared between two species. We paired DNA accessibility data with the gene expression data by separating chromatin accessibility clusters into three groups: early, late, and species-specific. The clusters in the early group were classified by the highest accessibility in the first half of each time-course while the late group was classified by the highest accessibility in the later stages of each time-course. The species-specific group is defined by differing accessibility trends between the rat and human in a cluster. There are 7,689 (41%) regions in six early clusters (JDA3, JDA4, JDA5, JDA9, JDA12, JDA13). Specifically, three of these clusters (JDA3, JDA9, JDA13) showed the early expression profiles in both rat and human. There are 4,506 (24%) regions in three late clusters (JNA11, JDA14, JDA15), and collectively show the late expression patterns in both species. In addition, we also found 6,747 (35%) regions in six species-specific clusters (JDA1, JDA2, JDA6, JDA7, JDA8, JDA10). Clusters in the early group included regions surrounding *POU5F1* (JDA3, JDA6, JDA9, JDA10, JDA13), *NANOG* (JDA13), *SOX2* (JDA3, JDA8, JDA9, JDA13), *MYCN* (JDA3, JDA6, JDA9, JDA13), and *TCF7L1* (JDA3, JDA4, JDA13, JDA15), which marked the stemness and early differentiation processes. Regions that are in the late clusters include *ID3* (JDA2, JDA7, JDA11, JDA13), *SOX17* (JDA4, JDA6, JDA11), *FOXA2* (JDA11), *FOXA1* (JDA4, JDA11, JDA14), *EOMES* (JDA1, JDA11, JDA15), and *GSC* (JDA11) which marked the final stage of DE differentiation in both species. Corresponding to the gene expression profiles (Figure 5A), we found one intermediate cluster, JDA1, reached to the highest accessible level at day 2 for human and day 4 for rat DE differentiation and also marked by the chromatin region activity around Brachyury T. Therefore, we treated this cluster as the mesendodermal cluster. We also found a set of TFs associated with cell cycle regulation and biosynthetic processes were enriched in species-specific clusters. These TFs included *ETS1* (JDA8, JDA9, JDA11), *NRSF/REST* (JDA8), *ID3* (JDA2, JDA7, JDA11, JDA13). Interestingly, we found CTCF (JDA6, JDA15), which is involved in gene regulation by mediating the formation of chromatin loops (Splinter et al. 2006; Stadhouders, Filion, and Graf 2019; Zhao et al. 2019), in cluster JDA6 that had early activation in rat but late activity in human (Figure 5E). Genes within this cluster were significantly enriched in cellular response to growth factor stimulus and apoptotic signaling pathway, indicating different roles of CTCF in mediating the formation of transcription units in two species. While 65% of alignable cCRE show conserved changes in chromatin accessibility between the two species, another 35% show species-specific behavior.

In order to further characterize the TF regions in each of the groups of clusters we analyzed the GO terms for the clusters in each group. The early grouping of accessible regions were significantly associated with regulation of neuron differentiation (p < 1×10^−25^), developmental growth (p < 1×10^−18^), cell morphogenesis (p < 1×10^−23^) and response to growth factor (p < 1×10^−17^) (Figure 5F), which indicates the regulation of early differentiation. Cluster JDA11,14,15 were highly enriched with endodermal markers that showed active chromatin accessibility at the end of DE differentiation for both species (Figure 5E). Regions in cluster JDA11 are associated with kidney (p < 1×10^−22^) and gland development (p < 1×10^−20^) and regions in cluster JDA14 are involved in urogenital (p < 1×10^−17^) and heart development (p < 1×10^−15^). In addition, regions in JDA15 are associated with wounding (p < 1×10^−16^) and muscle structure development (p < 1×10^−10^) (Figure 5G). As for the mesendodermal cluster, JDA1, regions are involved in mesenchyme development (p < 1×10^−20^), Wnt signaling pathway (p < 1×10^−14^) and pattern specification (p < 1×10^−18^), which correspond well with the intermediate cluster JDR6 in gene expression profiles (Figure 5A). Interestingly, for the species-specific clusters JDA2,6,7,8, regions are highly enriched in gene regulation processes in response to the external stimulus. For example, JDA7 is associated with mRNA processing (p < 1×10^−16^) and histone modification (p < 1×10^−15^) and JDA8 is associated with negative regulation of gene expression (p < 1×10^−17^) and Wnt signaling pathway (p < 1×10^−9^). Furthermore, JDA2 participates in tissue morphogenesis (p < 1×10^−18^) and JDA6 is involved in response to growth factor stimulus (p < 1×10^−15^) and apoptotic signaling pathway (p < 1×10^−15^) (Figure 5H).

These results indicate that there are groups of genes and regulatory elements with conserved transcriptional patterns across species. Chromatin accessibility is complementary with gene expression to identify conserved regulatory modules during DE differentiation in two species. However, species-specific patterns may reflect the difference in growth condition and following generation of transcripts during DE differentiation.

### Integrative analysis of NPC Differentiation

We then compared the relative conservation of gene expression and cCRE clusters between rat and human during NPC differentiation. We analyzed Early, Late, and species-specific RNA and chromatin clusters from Figures 4 to identify significant enrichments of shared genes and cCREs (Figure 6A; see methods). We found that 2,309 cCRE (out of 7,400 in Figure 4E) were within 20kb of 1,530 differentially expressed genes (out of 3,562 in Figure 4A). We detected seven sets of significantly enriched interactions, five of which were conserved, three with late RNA and late ATAC, one with early RNA and early ATAC, one with early RNA and late ATAC (discordant; Table S12). Overall, we found 175 (8%) cCREs for 145 genes in these five conserved clusters including a total 14 TF cCREs associated with 11 TFs. TFs in the early enriched NPC group include *GBX2*. TFs with shared late NPC affiliations include *ID1* (JNR8, JNA13), *JUNB* (JNR8, JNA13), *EGR1* (JNR8, JNA13), and *KLF16* (JNR17, JNA12) and are very strong candidates for conserved temporal regulation of differentiation in both species. The remaining two enriched associations showed species-specific differences. TFs in species-specific affiliated clusters include *PBX1* (JNR3, JNA3), *ATF3* (JNR3, JNA3), and *RARG* (JNR3, JNA3). These TFs may differ in the temporal aspects of their regulation during NPC differentiation in the two species.

**Figure 6:**
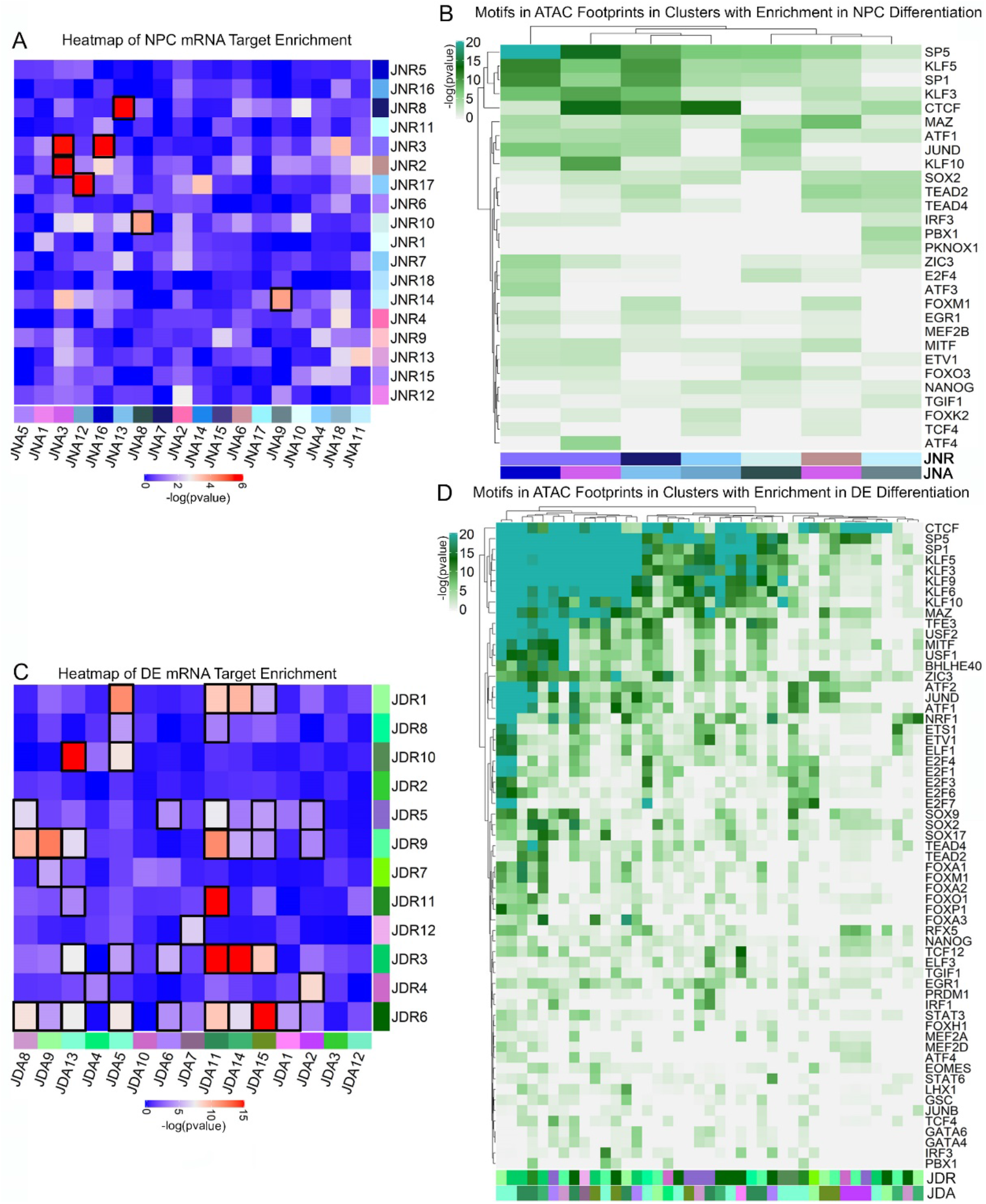
Integrative analysis of NPC and DE Differentiation. A) Heatmap of X^2^ p-values on a contingency table of clusters of accessible regions and expression clusters with applied P-value of 10^−4^ (Bonferroni corrected P-value: 0.05/[18 × 18]). The significant cluster overlaps are surrounded by a black box. The colors next to each cluster match the color bar from Figure 4A and 4E, and depict the affiliation with the early NPC group (light blue), late NPC group (dark blue), or species-specific group (pink). B) ATAC footprints in regions of significant overlap between clusters. The color bars are the same in Figure 4A, C, and E for each RNA and ATAC cluster. C) Heatmap of X^2^ p-values on a contingency table of clusters of accessible regions and expression clusters with applied P-value of 10^−4^ (Bonferroni corrected P-value: 0.05/[12 × 15]). The significant cluster overlaps are surrounded by a black box. The colors next to each cluster match the color bar from Figure 5A and 5E, and depict the affiliation with the early DE group (light green), late DE group (dark green), or species-specific group (pink). D) ATAC footprints in regions of significant overlap between clusters. The color bars are the same in Figure 5A, C, and E for each RNA and ATAC cluster.

We then mined the cCREs in the enriched clusters for footprints using Wellington and motifs using homer (Figure 6B). The most enriched footprints are SP5, KLF, SP1, and CTCF. These motifs are found in significant overlapping most clusters. Footprint enrichments for SOX2, IRF3, PBX1, and TGIF1 are mainly enriched in the Early cluster overlap and are potential candidate regulators for early NPC differentiation in both species. By contrast, footprint enrichments for SP1, KLF, ATF3, E2F4, EGR1, TCF4, TEAD, and ZIC3 are mainly found in Late cluster overlaps, which suggest that these TFs are likely candidate regulators for late NPC differentiation in both species. JUND and KLF10 footprint enrichments are found in the discordant cluster overlap. MAZ has footprint enrichment mainly in the species-specific cluster overlap, which suggests that MAZ may play a different role in NPC regulation in each species.

### Integrative analysis of DE Differentiation

We investigated the TFs that drive DE differentiation in both species using the same approach that we applied to NPC differentiation. We found that 10,148 cCRE (out of 18,942 in Figure 5E) were within 20kb of 5,322 differentially expressed genes (out of 7,320 in Figure 5A). We detected 41 sets of significantly enriched interactions between RNA and chromatin (see methods). 27 sets of interactions were conserved, including six with early RNA and early ATAC, seven with late RNA and late ATAC, seven with early RNA and late ATAC and seven with late RNA and early ATAC (Table S13). Overall, we found 2,214 (22%) cCREs for 1,548 genes in these conserved clusters, including a total 166 TF cCREs associated with 90 TFs. Some TFs with shared affiliations in early DE clusters include *SOX2* (JDR9, JDA9; JDR9, JDA13), *ETV1* (JDR1, JDA5), *IRF1* (JDR1, JDA5), *SOX11* (JNR8, JNA5), *FOXP1* (JDR9, JDA11), *GBX2* (JDR9, JDA13), *NANOG* (JDR9,JDA13), *CIR1* (JDR10, JDA5), and *LEF1* (JDR9, JDA13). Some of the TFs with shared late DE affiliations include *FOXA2* (JDR11, JDA11), *GATA4* (JDR11, JDA11), *EGR1* (JDR3, JDA11), *PITX2* (JDR11, JDA11), *JUND* (JDR3, JDA11), *TGIF1* (JDR3, JDA11), *FOXA1* (JDR6, JDA11), *ATF3* (JDR3, JDA11), *PRDM1* (JDR11, JDA11), and *TEAD2* (JNR6, JDA15). Some of the TFs with early RNA and late ATAC include *MYC* (JDR8, JDA11) and *GBX2* (JDR9, JDA14). Some of the TFs with late RNA and early ATAC include *TGIF1* (JDR3, JDA5), *T* (JDR6, JDA13), *JUN* (JDR3, JDA5), and *ZIC3* (JDR7, JDA9). These may be genes that are primed at different times before expression or are partially controlled by active repression by a repressive TF. The remaining 14 significantly enriched interactions are species-specific. TFs in species-specific affiliated clusters include *MEIS2* (JDR5, JDA8), *NRSF/REST* (JDR5, JDA8), *RARG* (JDR4, JDA2), *ID3* (JDR4, JDA2), *ETS1* (JDR5, JDA8, JDA11), *E2F1* (JDR4, JDA2), *TCF7L1* (JDR5, JDA15), *T* (JDR6, JDA1), and *FOXP1* (JDR9, JDA2). These TFs may differ in the temporal aspects of their regulation during DE differentiation in the two species.

We also mined the ATAC regions with significant overlap for footprints (Figure 6B). The most enriched footprints are CTCF, SP5, SP1, KLF, MAZ, and TFE3. These motifs are found in the majority of clusters and are general candidate regulators for both mesendoderm and DE differentiation of both species. Footprint enrichments for E2F4, E2F1, IRF3, E2F3, and E2F7 are mainly enriched in the conserved early clusters, making these TFs candidate regulators for early DE differentiation in both species. Footprint enrichments for JUND, LHX1, EGR1, bHLHE40, EOMES, TCF4, ZIC3, FOXA2/3, and TEAD are mainly enriched in late cluster overlaps. These TFs are strong candidate regulators for late DE differentiation in both species. ELF1, ETV1, GSC, JUNB, MEF2A, TCF12, TGIF1, and ATF3 motifs are found in clusters with significant overlapping clusters with discordant Early and Late affiliations. Footprint enrichments for GATA4/6, IRF1, NANOG, and PRDM1 are enriched in species-specific cluster overlaps, suggesting that these TFs are important candidate regulators for DE differentiation, but they differ in their temporal regulation in each species. The role of these TFs in DE differentiation are most likely not as evolutionary conserved as the TFs that share an early or late association between rat and human. Overall, we recovered many more significantly enriched interactions in DE differentiation than NPC differentiation.

## DISCUSSION

We used RNA and ATAC sequencing in a daily time-course to map gene and cCRE dynamics during differentiation into ectodermal neural progenitor cells and definitive endoderm in rat and human. The comprehensive gene expression and chromatin accessibility changes allowed us to identify conserved regulatory modules of lineage-specific regulators in both species. In particular, we were able to connect gene expression changes with associated changes of open chromatin regions during differentiation. We also found as expected that pluripotent gene expression was downregulated with the loss of chromatin accessibility. In addition, we identified substantial species-specific changes with instances of species-specific trajectories in rat and human during the same cell type differentiation. We found putative TF binding using open chromatin footprinting enriched in different sets of genes with conserved or species-specific changes. This is to our knowledge the first systematic comparison of *in vitro* differentiation from ESCs to ectodermal neural progenitor cells as well as to definitive endoderm in rat and human.

We find many cCRE similarities between NPC and DE in both species. For instance, the neuroectodermal marker, GBX2 is in significantly enriched early gene interactions in both NPC and DE time-courses in both species (Martinez-Barbera et al. 2001). EGR1 is found in significantly enriched late gene interactions in both NPC and DE time-courses in both species. Candidate regulators that increase in both species most likely have conserved function and temporal regulation. However, it is interesting to examine the genes that differed in expression over each time-course, as these genes may not be as conserved across rat and human. Genes like PBX1, ATF3, and RARG have species-specific expression and accessibility, suggesting they are required for different stages of NPC differentiation in the two species. Taken together, these candidate regulators are less evolutionarily conserved between species. Further experiments are required to understand the exact function and conservation of these candidate regulators in NPC differentiation between rat and human.

The conserved TFs FOXA2 and SOX17 were both up-regulated during DE differentiation, with relatively higher expression in rat than in human. CXCR4 is another key marker of mammalian endoderm differentiation (D’Amour et al. 2005; Mfopou et al. 2014; Morrison et al. 2016) and was upregulated in both species with higher expression in human. We also found that MIXL1 and EOMES starts expression early during DE differentiation, which has also been demonstrated in other organisms (Ryan et al. 1996; Lemaire et al. 1998; Colas et al. 2008). We also found that GATA4/6 and GSC are actively expressed in our time-course. Furthermore, we found that T, which is a marker of mesendoderm stage, was expressed highly in the middle of the time-course. This indicates that differentiating cells experienced the proper intermediate stage during endoderm development in both species. However, we found that some genes showed unexpected expression differences between species. The first is OCT4 (POU5F1), which is one of the key TFs defining the pluripotent cell fate in stem cells, that stayed highly expressed in later DE differentiation in human but not in rat. Previous studies have also found high expression of OCT4 at the end of human DE differentiation and they suggested this is possibly the effects of culture conditions (Lu et al. 2018). We also detected human-specific expression of NANOG in late DE stage that agrees with studies where OCT4 and NANOG induce DE markers during endoderm differentiation in human (Teo et al. 2011; Z. Wang et al. 2012). However, at least one endoderm single-cell study has shown the co-expression of OCT4 and endoderm markers in human DE (Chu et al. 2016). Further experiments are required to understand the exact function of OCT4 during DE differentiation as it may be one of the more important human-specific DE regulatory changes. In addition, we also detected expression of FOXA1 in rat but not in human during DE development. As a core member TF of the FOXA family, expression of FOXA1 and 2 have been detected in both mouse and Xenopus endoderm. Although a study showed that FOXA1 is the direct target of EOMES in human DE and knockdown EOMES would decrease the FOXA1 expression (Teo et al. 2011), our results indicate that FOXA1 may not be not an essential TF during human DE development as the equivalent function may be taken over by FOXA2.

Of the most enriched footprints shared between both NPC and DE differentiations SP5, KLF, SP1, and CTCF are needed in many cell types and for a broad range of cell functions (Arzate-Mejía, Recillas-Targa, and Corces 2018; Kaczynski, Cook, and Urrutia 2003). SP5 has been found to interact with brachyury/T during development (Harrison et al. 2000). Footprints for IRF3 are found in early significant cluster affiliations for both NPC and DE differentiations. This may be due to the exiting of the pluripotent state and into a more differentiated state, as pluripotent ESCs do not utilize IRF3 (Eggenberger et al. 2019) and it is a regulator of cell proliferation. Footprints for SOX2 are found in early NPC significant cluster overlaps due to its ability to regulate both pluripotent ESCs and multipotent NPCs (Lodato et al. 2013). Footprints for TGIF1 and PBX1 were also found in early NPC cluster overlaps. TGIF1 (van de Leemput et al. 2014) and PKNOX, a co-factor of PBX1, regulate neural cell fate specification (Golonzhka et al. 2015). Footprints for TEAD, EGR1, TCF4, and ZIC3 are found in late significant cluster affiliations for both NPC and DE differentiations. ZIC3 and EGR1 regulate the transition out of naïve pluripotency (Yang et al. 2019; Kalkan et al. 2017) and TCF4 has been shown to regulate cell survival and neuronal differentiation (Forrest et al. 2013). Whereas members of the TEAD family are needed for the regulation of differentiation into both endoderm and neuronal lineages (Mukhtar et al. 2020; Cebola et al. 2015). Footprint enrichments for MAZ are also shared in NPC and DE differentiations. MAZ is part of the most enriched footprints in DE differentiation whereas it shows species-specific enrichment in NPC differentiation. MAZ has been shown to mediate promoter activity during neuronal differentiation (Okamoto et al. 2002) and SP1 and MAZ may share the same cis-regulatory elements (Song et al. 2001). Further experiments are required to understand the exact function and conservation of these candidate regulators in NPC and DE differentiation.

One of the most striking results of our comparative analysis of NPC and DE differentiation is that we find considerably more conservation during DE differentiation than NPC differentiation. This could be a product of the *in vitro* system or inherent biological differences. One possible reason is the differences between differentiation protocols for each species. Both NPC differentiation protocols concluded after eight days but used different inhibitors. For human NPC differentiation, three inhibitors were added to the media (ALK2/3, ALK5, and GSK3) while only one was used in rat (ALK2/3) with the addition of the growth factor FGF2. For DE differentiation, the 5-day human protocol was performed using an established kit while in rat the 7-day rat protocol used a plethora of growth factors and inhibitors such as Activin. The differentiation of both these cell-types took equally as long or longer in rat, which is a key difference from *in vivo* where the formation of both these cell-types occurs on a faster scale in rat (Guillaume and Zhang 2008; Murry and Keller 2008). Variations in differentiation efficiency for each cell-type between the two species is also a possibility. Another class of differences could be biological. For example, our results could be impacted by different starting ESC states between rat and human. Human H1 ESCs are in a primed state (post-implantation; epiblast cells) while mouse ESCs are in a naive state (preimplantation blastocyst; inner cell mass) (Takahashi, Kobayashi, and Hiratani 2018). The rat DAc8 ES cells are thought to be in-between the naive state of the mouse ESCs and primed state of the H1 ESCs. Rat ESCs are cultured in similar conditions to naive mouse ESCs, with medium containing two inhibitors (2i), but express Cdx2, which is a trophoblast marker not expressed in naive mouse ESCs (Hong et al. 2013). Another possible explanation is the differing endpoints between DE and NPC differentiation. The complexity of neuroectoderm and NPC differentiation *in vivo* is extremely difficult to recapitulate *in vitro*. These cells could have positional identity *in vivo* such that their fate is influenced by environmental cues from neighboring cells, which can only be partially recapitulated *in vitro*. Rodents and humans have a large gestational time gap in neuroepithelial cell formation. In rodents these cells can be distinguished after about 7 days gestation but are only distinguishable near the end of the third week of gestation in humans. Numerous NPC subtypes exist, many of which can be recapitulated *in vitro* (Guillaume and Zhang 2008). Formation of DE *in vivo* occurs much earlier in development than the formation of NPCs. DE develops from epiblast cells that transition from the most anterior region of the primitive streak, which marks the beginning of gastrulation (Murry and Keller 2008). Previous studies have shown that there is a common pathway for inducing definitive endoderm formation that is shared between many species (Yiangou et al. 2018). This similarity of early development of DE between species paired with the complexity of later formation of NPCs and developmental time differences between species may account for the striking differences in conservation seen between NPC and DE differentiations.

We used transcriptome and cCRE changes during the differentiation of rat and human ESCs to map expression changes and cCRE dynamics during differentiation into ectodermal neural progenitor cells and definitive endoderm lineages. Our gene expression time-courses allowed us to carefully define temporal profiles of expression in connection with the timing of early differentiation in the two germ layers. We can discover evolutionarily conserved and species specific *cis*-regulatory elements by comparing conserved DNA between evolutionarily rodents and humans whose lineages diverged about 91 million years ago (F. G. Jørgensen et al. 2005). While mice are often used in comparative genomic analyses with human (Stergachis et al. 2014), rats have been less intensively studied using functional genomics for comparative purposes. In particular, mice and rats diverged around 12 million years ago, there is likely to be very interesting and novel evolutionary changes found between rat and human that are not found in mouse (Ramsdell et al. 2008), (Hardison and Taylor 2012). A comparative analysis of GRNs between the three species would be an important next step to understand how the changes that we see in RNA and ATAC during differentiation can occur while conserving the overall process of DE and NPC differentiation.

## AUTHOR CONTRIBUTIONS

A.M., S.J., and C.W.T. conceived the study. C.W.T. performed ESC culture and NPC differentiation, RNA-seq and ATAC-seq libraries preparation, as well as analysis of RNA-seq and ATAC-seq data, and footprinting analysis; S.J. performed DE differentiation, RNA-seq and ATAC-seq libraries preparation, and data analysis. C.M. and S.M. sequenced the RNA-seq and ATAC-seq libraries. C.W.T, S.J., and A.M. wrote the manuscript.

## METHODS

### Embryonic stem cell maintenance

H1 (male) human embryonic stem cells (ESCs) were obtained from WiCell and maintained in STEMCELL Technologies TeSR-E8 medium on growth factor reduced (GFR) Corning Matrigel. Cells were routinely passaged every 2-3 days with 0.5mM EDTA in dPBS. DAc8 (male) rat embryonic stem cells were purchased from Rat Resource and Research Center (RRRC), University of Missouri. Cells were first maintained on mouse embryonic fibroblast (MEF) feeders plated on 0.1% gelatin. MEF medium was made with GMEM, 10% FBS,1% GlutaMAX and 1% Pen/Strep. Rat ES cells were cultured in a 1:1 mixture of DMEM/F12 and Neurobasal medium supplemented with N2, HEPES (1M), B27, GlutaMax-I, Insulin, CHIR99021 (3mM), PD0325901 (1mM), 2ß-ME, Y-27632 (5mM), and hLIF(10ug/ml). MEF cells were plated at least one day before plating rat ES cells. Rat ES cells were passaged every 4-6 days with Accutase.

### Definitive endoderm (DE) differentiation on monolayer *in vitro*

Human ES cells were differentiated using the STEMdiff Definitive Endoderm Kit (TeSR-E8 Optimized) from STEMCELL Technologies (Ramme et al. 2019). Rat ES cells were differentiated following an optimized mouse protocols (Kim et al. 2010; Yasunaga et al. 2005; Mfopou et al. 2014). For rat, cells were first transferred from MEFs to gelatin coated plate and they were seeded in rat ES medium for 4-6 hours.Then the medium was changed to the NDiff N2B27 basal medium supplemented with PD0325901 (1uM), CHIR99021 (3uM) and mLIF (100U/ml) (in 50ml, 49.939 ml of NDiff, 1ul mLIF, 10ul PD03, 50ul CHIR) on 0.1% gelatin. After 2-3 days, medium was changed to NDiff medium supplemented with Activin A, Fgf4, Heparin, kinase inhibitor (PI103), and CHIR99021 (in 50ml, 49.88ml of NDiff medium, 10ul of 100ug/ml ActivinA, 5ul of 100ug/ml Fgf4, 50ul of 1mg/ml of Heparin, 5ul of 1mM PI103, 50ul of 3uM CHIR). After 2 days of differentiation, medium was changed to DMEM/F12 with N2, B27-VA, L-glutamine, 2ß-ME, BSA, ActivinA, Fgf4, Heparin, Egf, PI103 and CHIR99021 (in 50ml,48.8295ml of DMEM/F12,250ul of N2, 500ul of B27-VA, 250uL-glutamine, 50ul 1000x 2ß-ME, 0.025g BSA, 10ul 100ug/ml ActivinA, 5ul 100ug/ml Fgf4, 50ul 1mg/ml Heparin, 0.5ul100ug/ml Egf, 5ul 1mM PI103 and 50ul of 3uM CHIR) for 5 days.

### Neural Progenitor Cell (NPC) Differentiation on monolayer *in vitro*

Human ES cells were differentiated using an adapted previously established protocol (Porterfield 2020). Briefly, cells were passaged and plated on matrigel and allowed to reach 30-40% confluence. After which, media was changed to NPC medium (1:1 IMDM/F12 supplemented with NEAA, N2, B27, PSA, ALK2 and ALK3 inhibitor [0.2uM LDN193189], ALK5 inhibitor [10uM SB431542], and GSK3 Inhibitor [3uM CHIR99021]) for 8 days. Rat ES cells were also differentiated using a modified previously published protocol (Alsanie et al. 2017). For rat, cells were first transferred from MEFs to gelatin coated plate and they were seeded in rat ES medium for 4-6 hours before changing to 1:1 IMDM/F12 differentiation medium supplemented with NEAA, PSA, N2, B27, LDN193189 (200 nM, Sigma-Aldrich), with FGF2 (20 ng/ml, Sigma-Aldrich) added to the media from day 2-5.

### RNA-seq library construction

Total RNA was extracted once a day during differentiation using the RNeasy kit (QIAGEN). RNA was converted to cDNA using the SmartSeq2 protocol (Picelli et al. 2014). Libraries were constructed by using the Nextera DNA Flex Library Prep or Illumina DNA Prep kit (Illumina). Libraries were quality-controlled prior to sequencing using Agilent 2100 Bioanalyzer profiles and normalized using the KAPA Library Quantification Kit (Illumina). The libraries were sequenced using Illumina NextSeq500 platform and sequenced to a depth of around 10 million reads per sample.

### ATAC-seq library construction

ATAC-seq samples were collected from the same pool cells collected daily for RNA-seq and flash frozen based on the omni-ATAC protocol (Corces et al. 2017). Around 75,000 cells were used per replicate and libraries were size-selected between 150-500bp using electrophoresis as done previously (Ramirez et al. 2017). In short, gel was viewed and cut between 150-500bp and DNA was extracted from gel pieces using QIAquick Gel Extraction Kit (QIAGEN). Libraries were normalized using the KAPA Library Quantification Kit (Illumina). The libraries were sequenced using Illumina NextSeq500 platform and sequenced to a depth of around 60-100million reads per sample.

### Gene expression analysis

Raw reads were mapped to hg38 (human), and rn6 (rat) using STAR (version2.5.1b) (Dobin et al. 2013) using default except with a maximum of 10 mis matches per pair, a ratio of mis matches to read length of 0.07, and a maximum of 10 multiple alignments. Quantitation was performed using RSEM (version1.2.31) (Li and Dewey 2011) with the defaults, and results were output in transcripts per million (TPM) and counts. Batch effects between species were corrected by using limma removebatcheffect (Smyth, n.d.). Clustering of differentially expressed genes across the time-course was done by using maSigPro (Conesa et al. 2006) with alpha of 0.05 for multiple hypothesis testing and a false discovery rate of 0.05%. Gene ontology analysis was done by using Metascape (Zhou et al. 2019).

### ATAC-seq data processing and analysis

Raw reads were mapped to hg38 (human) and rn6 (rat) using bowtie2 (Li and Dewey 2011; Langmead and Salzberg 2012).Reads mapped to ChrM were discarded and PCR duplicates were removed by using Picard (“Website” n.d.). HOMER/4.7 (Heinz et al. 2010) was used to call candidate regulatory elements (cRE). It was first used for calling 150bp narrow peaks and then 500bp broad peaks. Then narrow and broad peaks were merged into a single peak list and they were further filtered by overlapping with ENCODE “blacklist” regions. Peaks that were shown in both replicates were considered as reproducible cRE. Reads coverage was calculated using bedtools (Quinlan and Hall 2010) for each region and they were further normalized by the size of the library and the size of peaks. Normalized coverages in inter-species comparisons were further batch corrected for the technical batch effects by using limma removebatcheffect (Smyth, n.d.). The differential cCRE regions through differentiation time-course were identified by using maSigPro (Conesa et al. 2006) with alpha=0.05 and FDR<0.05.4.6.7 de novo motif enrichment analysis denovo motif calling was performed using HOMER/4.7 (Heinz et al. 2010) using masked genomes for each species. The size of motif was set as “-len 6, 8, 10, 12, 15, 120” with at most 3 mismatches.

### Contingency Tables and X2 test

RNA and ATAC gene overlaps were counted for each cluster pair and a χ2 test was used to determine significance for each cluster overlap. To study the enrichment of accessibility regions in different mRNA clusters for Figure 2E and Figure 3E an applied P-value of 10^−4^ (Bonferroni corrected P-value: 0.05/[18 × 24]) was used (Table S4 and S7). Figure 6A overlaps had an applied P-value of 10^−4^ (Bonferroni corrected P-value: 0.05/[18 × 18]; Table S12) and Figure 6C overlaps had an applied P-value of 10^−4^ (Bonferroni corrected P-value: 0.05/[12 × 15]; Table S13).

### Footprint Calling and GRN Construction

ATAC-seq reads from duplicates were merged to achieve at least 100-120 million reads for footprints calling. Footprints were called in chromatin regions from clusters with significant overlap with mRNA clusters by using Wellington algorithm from pyDNase (Piper et al. 2014) with parameters “-A -fp 6, 31, 1 -sh 7, 36, 4 -fdr limit -2”, 1%FDR. Then motifs were scanned within footprints regions by using homer (Heinz et al. 2010).

### Inter-species analysis

Inter-species pairwise comparisons were performed by aligned identified cCRE between species in a reciprocal manner using UCSC liftOver (Kent 2002) on genomic assemblies in two species. Each of the species was used as an anchor species and the cCRE were mapped to the other 2 species with 50% minimum map ratio. Regions failing to be mapped in any of the other genomes were considered as unaligned regions. To identify conserved regions between two species, regions having orthologous regions overlapped in the second species with at least 1 bp were collected. Final pair wise conserved regions were confirmed by doing this comparison reciprocally between species

## Acknowledgments

We thank Claire Zeng and Katherine Williams for feedback on the project and manuscript. This work was supported in part by grants from the National Institutes of Health DP2 GM111100 and UM1 HG009443 to A.M

## Data and Software Availability

The accession number for the sequencing data reported in this paper is GEO: GSE164361

## Supplemental Figures

Figure S1: Related to Figure 1. **Monolayer differentiation of embryonic stem cells into definitive endoderm and neural progenitor cells**

Figure S2: Related to Figure 2, **correlation and visualization of differential ATAC and RNA profiles over rat NPC and DE differentiation time-courses**

Figure S3: Related to Figure 2, **gene ontology and motif analysis of differential open chromatin regions in clusters**

Figure S4: Related to Figure 3, **correlation and visualization of differential ATAC and RNA profiles over human NPC and DE differentiation time-courses**

Figure S5: Related to Figure 3, **gene ontology and motif analysis of differential open chromatin regions in clusters**

## Supplementary tables

Table S1: **Statistics for RNA and ATAC experiments**

Table S2: **Cluster analysis of differential gene expression between NPC and DE differentiation in rat**

Table S3: **Cluster analysis of differential chromatin accessibility between NPC and DE differentiation in rat**

Table S4: **Correlation of mRNA-ATAC clusters in rat NPC and DE comparison analysis**

Table S5: **Cluster analysis of differential gene expression between NPC and DE differentiation in human**

Table S6: **Cluster analysis of differential chromatin accessibility between NPC and DE differentiation in human**

Table S7: **Correlation of mRNA-ATAC clusters in human NPC and DE comparison analysis**

Table S8: **Cluster analysis of differential gene expression between rat and human NPC differentiation**

Table S9: **Cluster analysis of differential chromatin accessibility between rat and human NPC differentiation**

Table S10: **Cluster analysis of differential gene expression between rat and human DE differentiation**

Table S11: **Cluster analysis of differential chromatin accessibility between rat and human DE differentiation**

Table S12: **Correlation of mRNA-ATAC clusters in rat and human NPC differentiation**

Table S13: **Correlation of mRNA-ATAC clusters in rat and human DE differentiation**

