## Supplemental Figures for "Comparative Chromatin Dynamics of Stem Cell Differentiation in Human and Rat"

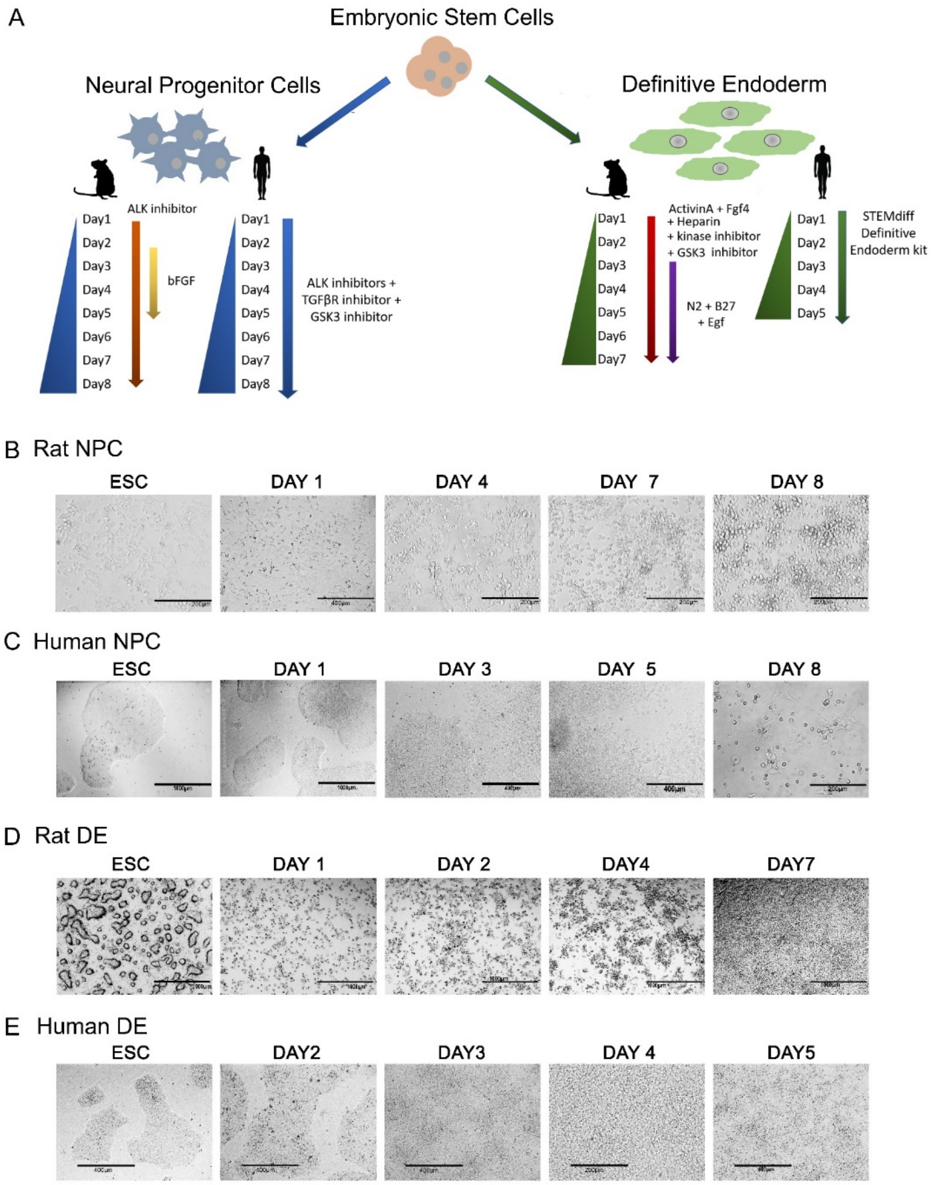


Figure S1: Related to Figure 1. **Monolayer differentiation of embryonic stem cells into definitive endoderm and neural progenitor cells**

A) Schematic representation of study design. Color indicates differentiation cell lineage. Human or rat symbols denote species. Arrows indicate media composition.

B) Brightfield images of eight day time-course of rat neural progenitor cells by using adapted protocol (Alsanie et al. 2017)

C) Brightfield images of eight day time-course of human neural progenitor cells by using adapted protocol (Porterfield 2020).

D) Brightfield images of seven day time-course of rat definitive endoderm differentiation by using adapted protocol (Mfopou et al. 2014)

E) Brightfield images during five day time-course of human definitive endoderm differentiation by using STEMdiff Definitive endoderm kit

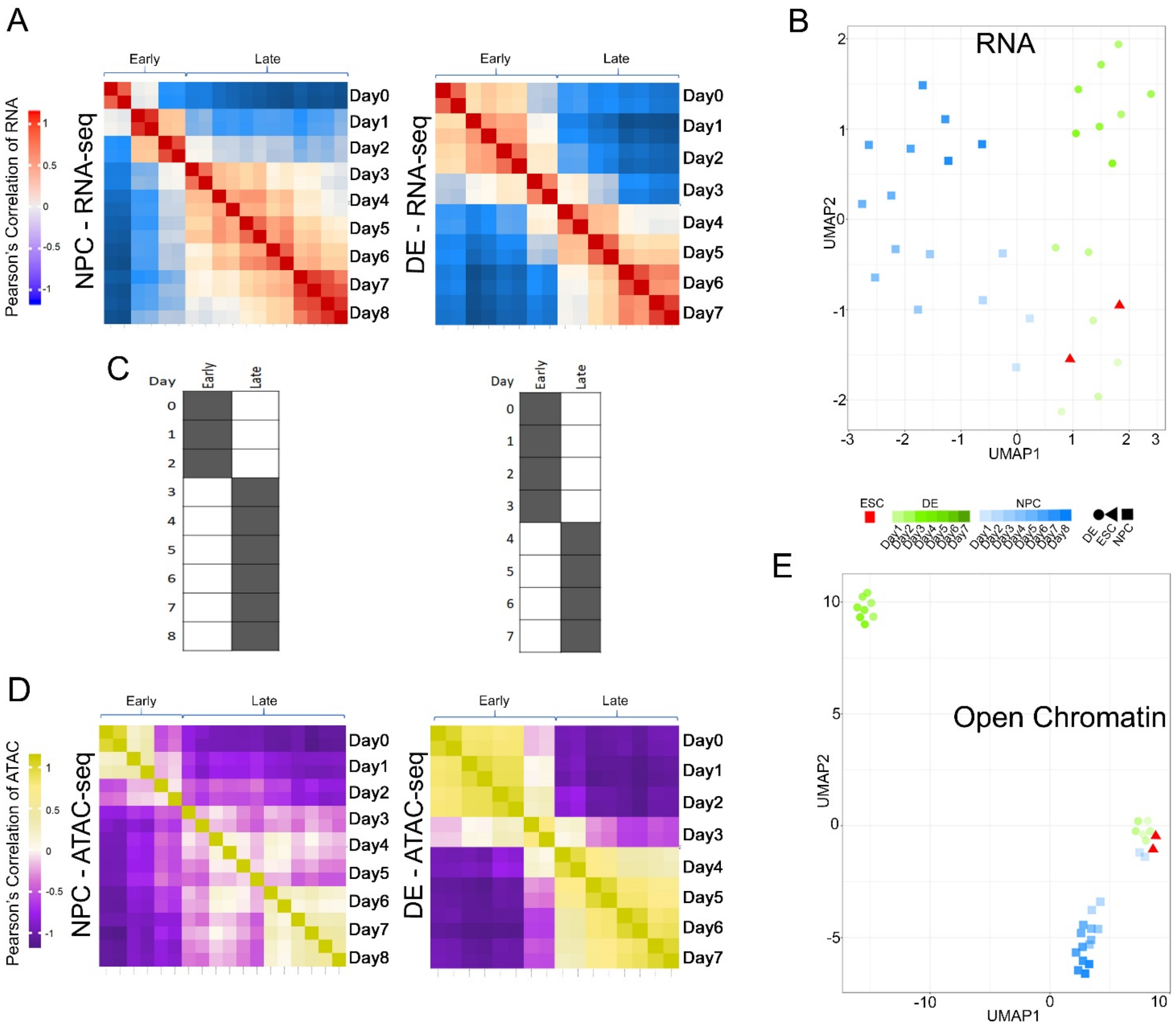


Figure S2: Related to Figure 2, **correlation and visualization of differential ATAC and RNA profiles over rat NPC and DE differentiation time-courses**

A) Pearson correlation of expression profiles over rat NPC and DE differentiation time-courses. The samples are separated into early and late expression profiles for each time-course.

B) UMAP representation of differential expression profiles of rat NPC and DE time-courses annotated by cell-types. Color darkens as time-course progresses.

C) Temporal staging of time points for each cell type was based on the clustering of RNA-seq and ATAC-seq data.

D) Pearson correlation of open chromatin profiles over rat NPC and DE differentiation time-courses. The samples are separated into early and late expression profiles for each time-course.

E) UMAP representation of differential open chromatin profiles of rat NPC and DE time-courses annotated by cell-types

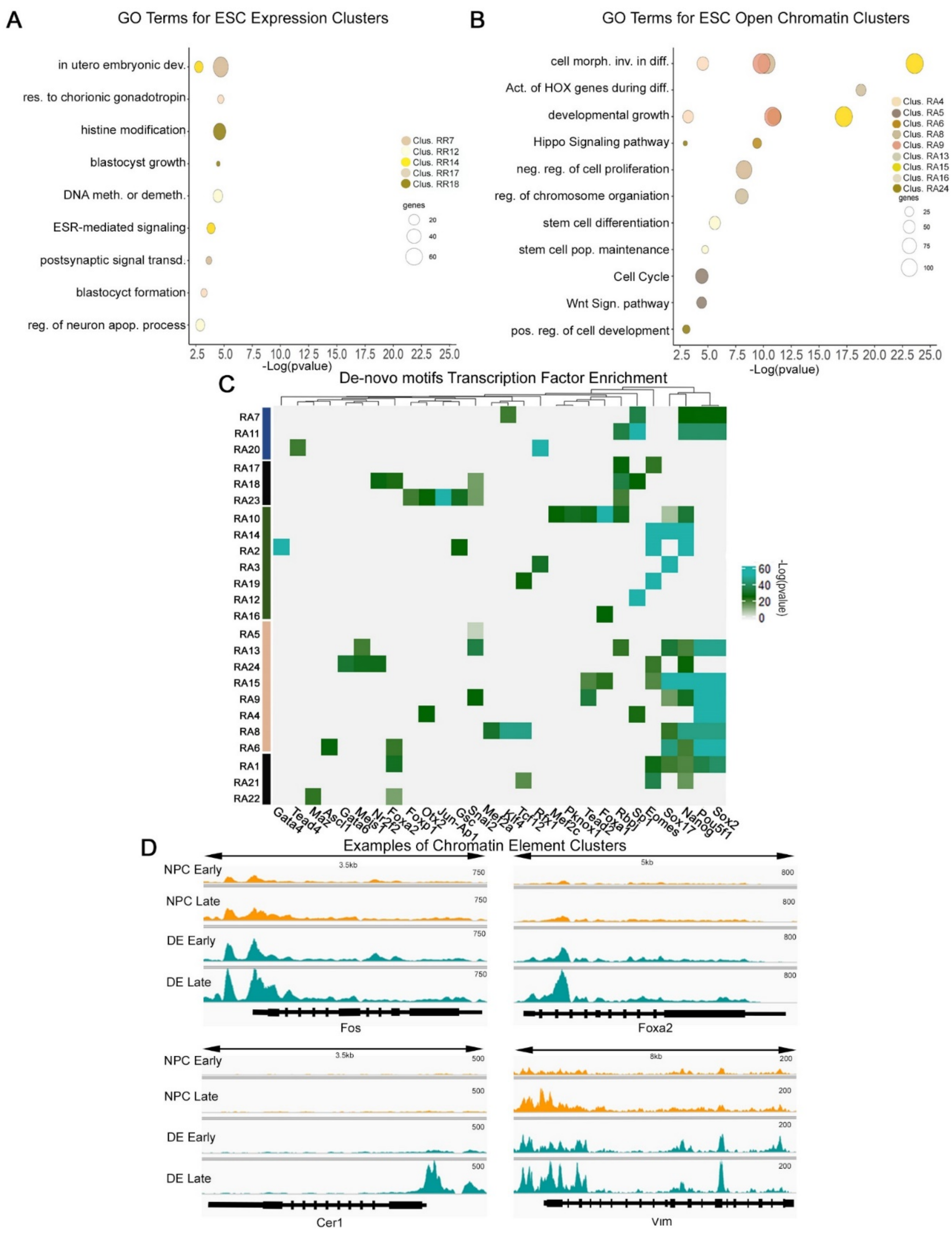


Figure S3: Related to Figure 2, **gene ontology and motif analysis of differential open chromatin regions in clusters**

A) Gene ontology (GO) enrichment analyses of differentially expressed rat genes in ESC category

B) GO enrichment analyses of differential open chromatin rat regions in ESC category

C) Motif enrichment between differential open chromatin regions in clusters. Clusters on the left are in the same order as the differential heatmap in Figure 2C

D) Examples of four open chromatin elements in rat visualized in Integrative Genome Viewer (IGV) that increase in either NPC (orange) or DE (teal) differentiation. Mapped ATAC reads are separated into early and late groupings from Figure S2E.

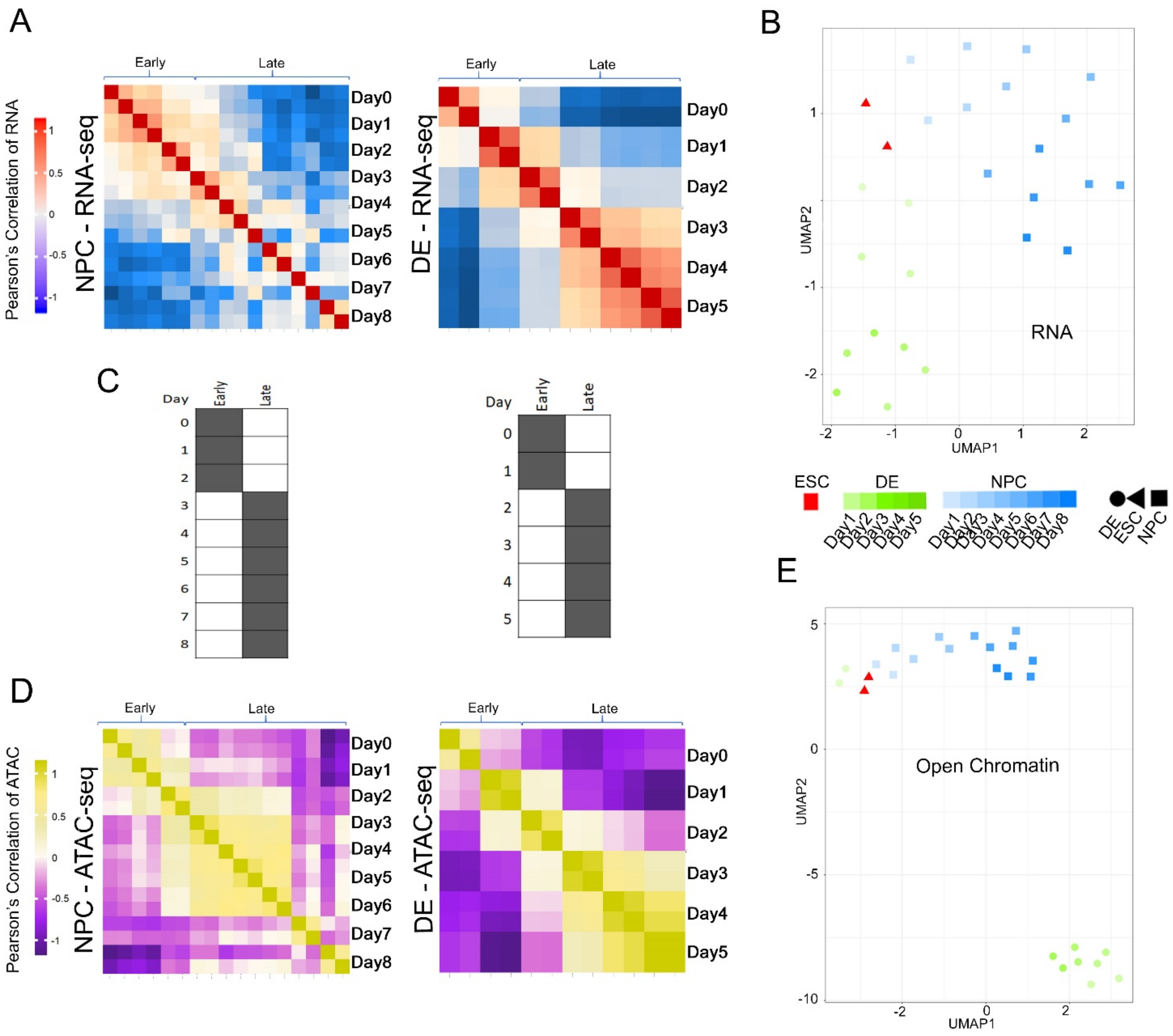


Figure S4: Related to Figure 3, **correlation and visualization of differential ATAC and RNA profiles over human NPC and DE differentiation time-courses**

F) Pearson correlation of expression profiles over humanNPC and DE differentiation time-courses. The samples are separated into early and late expression profiles for each time-course.

G) UMAP representation of differential expression profiles of human at NPC and DE time-courses annotated by cell-types. Color darkens as time-course progresses.

H) Temporal staging of time points for each cell type was based on the clustering of RNA-seq and ATAC-seq data.

I) Pearson correlation of open chromatin profiles over human NPC and DE differentiation time-courses. The samples are separated into early and late expression profiles for each time-course.

J) UMAP representation of differential open chromatin profiles of human NPC and DE time-courses annotated by cell-types

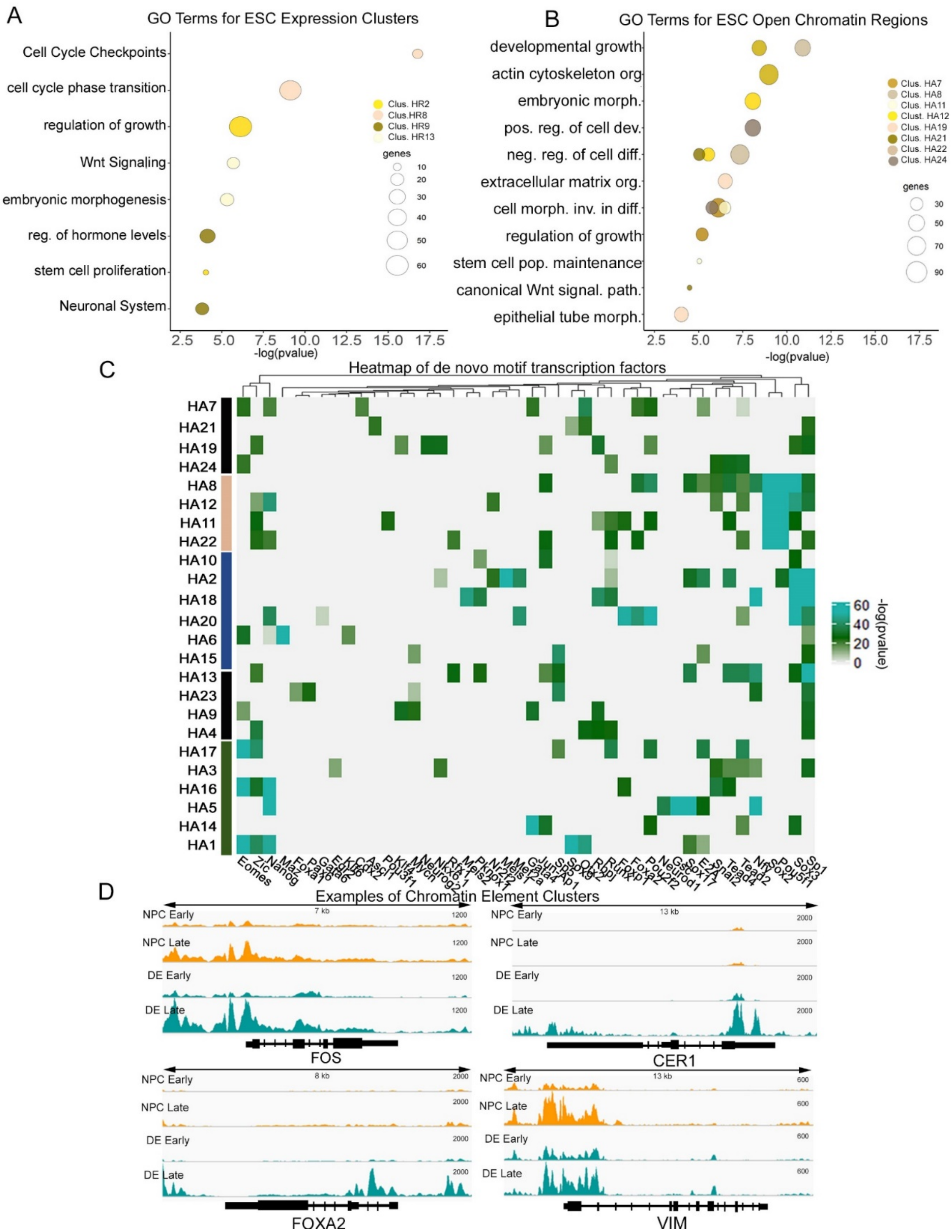


Figure S5: Related to Figure 3, **gene ontology and motif analysis of differential open chromatin regions in clusters**

E) GO enrichment analyses of differentially expressed human genes in ESC category

F) GO enrichment analyses of differential open chromatin human regions in ESC category

G) Motif enrichment between differential open chromatin regions in clusters. Clusters on the left are in the same order as the differential heatmap in Figure 3C

H) Examples of four human open chromatin elements visualized in Integrative Genome Viewer (IGV) that increase in either NPC (orange) or DE (teal) differentiation. Mapped ATAC reads are separated into early and late groupings from Figure S4E.
